# Environmental anchoring of grid-like representations minimizes spatial uncertainty during navigation

**DOI:** 10.1101/166306

**Authors:** Tobias Navarro Schröder, Benjamin W. Towse, Matthias Nau, Neil Burgess, Caswell Barry, Christian F. Doeller

## Abstract

Minimizing spatial uncertainty is essential for navigation, but the neural mechanisms remain elusive. Here we combine predictions of a simulated grid cell system with behavioural and fMRI measures in humans during virtual navigation. First, we showed that polarising cues produce anisotropy in motion parallax. Secondly, we simulated entorhinal grid cells in an environment with anisotropic information and found that self-location is decoded best when grid-patterns are aligned with the axis of greatest information. Thirdly, when exposing human participants to polarised virtual reality environments, we found that navigation performance is anisotropic, in line with the use of parallax. Eye movements showed that participants preferentially viewed polarising cues, which correlated with navigation performance. Finally, using fMRI we found that the orientation of grid-cell-like representations in entorhinal cortex anchored to the environmental axis of greatest parallax information, orthogonal to the polarisation axis. In sum, we demonstrate a crucial role of the entorhinal grid system in reducing uncertainty in representations of self-location and find evidence for adaptive spatial computations underlying entorhinal representations in service of optimal navigation.

## INTRODUCTION

Accurate navigation is a daily challenge for humans and other animals alike. In order to self-localise an agent must integrate incomplete and uncertain information regarding its position and motion. In the brain, entorhinal grid cells are thought to play a central role in this process^1–6^ – appearing to provide an efficient representation of self-location updated on the basis of self-motion and the proximity to salient cues. However, while sensory representations - such as those found in the visual cortex - are known to adapt in response to varying levels of uncertainty^7, 8^, it is unknown whether spatial representations in the medial temporal lobes respond similarly.

In rodents, the regular triangular firing-patterns of grid cells can be heavily distorted by environmental geometry - tending to align to walls of rectangular enclosures^9, 10^. It becomes fragmented in hair-pin mazes and distorted in trapezoidal environments^10, 11^, which translates into systematic memory distortions in humans^12^. Human entorhinal fMRI- and iEEG activity during virtual navigation is modulated by movement direction. This modulation shows six-fold rotational (hexadirectional) symmetry that has been proposed as a population signal of grid cells (i.e. grid-cell-like representations)^13–20^. These grid-cell-like representations are present also during visual exploration and anchor to square boundaries^16, 17^. Similar activity patterns have been observed in monkeys during visual tasks^21^.

Currently, it is unclear whether these distortions are maladaptive - a failed attempt to generate a regular grid, which might conceivably result in navigational errors^22^. However, an alternative explanation could be that the irregularities confer an advantage, supporting more accurate self-localisation than regular grid-patterns would. Indeed, theoretical considerations suggest that the transient expansion in grid scale observed when animals are exposed to novel enclosures^23^ may be a strategy to minimise decoding errors in unfamiliar and hence uncertain environments^2^. Plausibly similar adaptive processes would generate altered grid-patterns in response to asymmetries or local variability in the availability of reliable spatial cues.

Here, we test if polarised spatial cues – providing anisotropic motion parallax information during self-motion - systematically alter the configuration of grid-like representations in a way that is consistent with the minimisation of decoding errors. To this end, we employed a simulated grid cell system to make specific predictions about the optimal orientation of grid-patterns under conditions of asymmetric spatial information. In turn, we tested these predictions against the orientation of human grid-cell-like representations monitored while human subjects performed a virtual navigation task. Eye-tracking data and behavioural performance measures were used to assess the participants’ use of spatial cues and their ability to navigate accurately within the VR.

## RESULTS

### Impact of environmental geometry on grid pattern may be adaptive

Computational simulations suggest that grid cell firing patterns can partially mitigate the effects of increased spatial uncertainty by increasing in scale^2^ – an effect observed empirically in novel environments ^23^. We build on this existing framework ^2, 4^, inquiring how position decoding using a population of grid cells with coherent orientation is affected by anisotropy in spatial uncertainty (random displacement of each grid pattern along a cardinal axis). Grid cell ensembles with an orientation offset at 0 to 30° relative to the axis of greater uncertainty were simulated (in a six-fold rotational system 30° is the greatest possible offset).

The simulations (see Materials and Methods) demonstrated a substantial difference in the accuracy with which self-location can be decoded depending on the orientation of the grid pattern relative to the axis of greatest uncertainty. Specifically, the most accurate representation of position was obtained when the grid population was oriented at 30° to the axis - equivalent to being fully misaligned to the axis defined by the spatial cues; Figure 1C, Figure 1 – figure supplement 1-3). Decreasing the anisotropy in spatial uncertainty diminished this effect, until no directional benefit was apparent for isotropic uncertainty – a spherical noise distributions (Figure 1 and Figure 1 – figure supplement 1). In further control experiments, we varied grid scale, firing rate, and the number of grid modules – the optimal grid orientation relative to an axis of uncertainty remained consistent (Figure 1 – figure supplement 2). Similarly, in the case of less plausible, higher-order noise distributions with multiple peaks, an optimal grid orientation was only present if the alignment of the grid pattern and the noise distribution could be reduced through rotation. This was the case for a six-leaf distribution, but not for a four-leaf distribution (Figure 1 – figure supplement 3).

**Figure 1.**
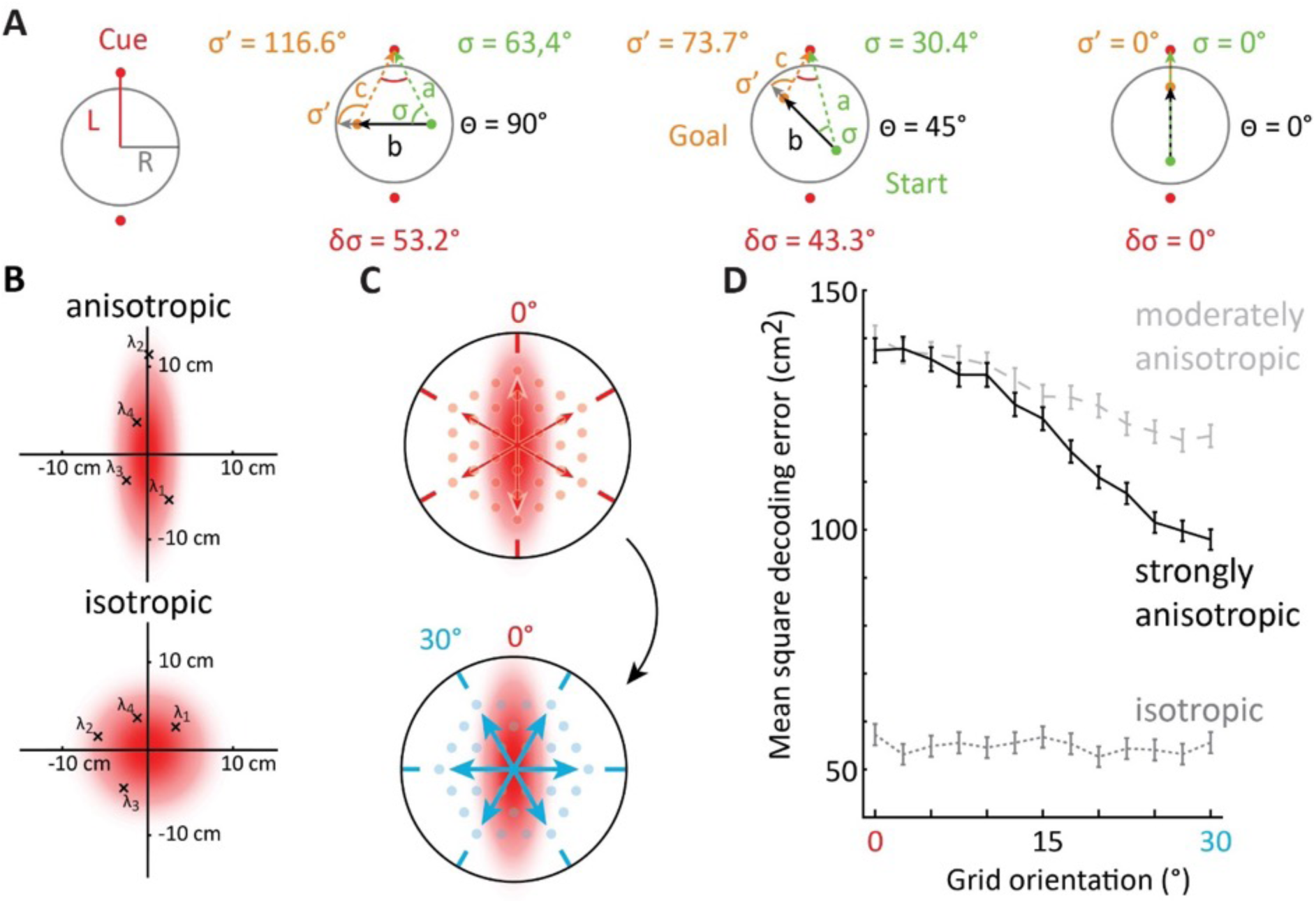
Spatial information is anisotropic and affects optimal self-localisation using grid cells. **A** Spatial information during movement in polarised environments is anisotropic. Left: schematic of arena with radius *R*, and polarising cues at distance *L.* Right: three example paths in different directions and through the centre are shown (black arrows at angle θ; Path lengths are equal to distance of the cue to the centre). The change in cue direction from the observer’s heading (δσ) is maximal on paths perpendicular to the polarisation axis (note: this is also the case on average if paths are not centred on the middle but distributed evenly throughout the environment). **B-D** We simulated decoding of position estimates from the activity of grid cell ensembles with patterns oriented at different angles relative to the axis of lowest spatial certainty. Note that the axis of lowest spatial certainty in A is the polarisation axis formed by the two cues, because angular change is smallest and triangulation errors are largest along it. Uncertainty in spatial information for the simulations of position decoding using grid cells was introduced by adding Gaussian errors to the true position input. These errors were generated independently for each module of grid cells. Anisotropy was created by separately varying the standard deviations of the error in two orthogonal axes. **B** illustrates an example: the subject’s actual location is at the origin; red shading indicates a two-dimensional probability density distribution for error generation, with either different or equal standard deviations in each axis (anisotropic and isotropic, respectively); and crosses indicate four independently generated noisy position estimates, drawn from this distribution and be input to each of the grid cell system’s four modules. **C** Schematic illustration of two grid orientations either aligned with the uncertainty axis (left panel, arrows indicate hexadirectional orientations associated with a grid), or rotated 30° (right panel). The number of depicted grid fields differ only for illustration purpose. **D** Position decoding error, defined as the mean maximum-likelihood estimate square error (MMLE; cm^2^), was largest when one grid axis was aligned at 0° relative to the axis of lowest spatial certainty (as shown in panel B, top). No optimal grid orientation was present in the isotropic condition. Solid black line: ‘strongly anisotropic’, errors with s.d. 5cm and 0cm; dashed light grey line: ‘moderately anisotropic’, 5cm and 1.67cm; dotted medium grey line: ‘isotropic’, 3.33cm in all directions. Grid orientation is defined as the minimal angular offset of a grid axis from the axis of greater uncertainty (this is analogous to hexadirectional offset of entorhinal fMRI activity described below). Error bars indicate 95% confidence interval (n=150,000).

### Motion parallax is maximal during movement perpendicular to polarising cues

To investigate the effects of uncertainty on grid cells and navigation, we first sought to generate a virtual environment characterised by anisotropic spatial information. Because motion parallax - the apparent change in bearing to stationary points - is a major source of spatial information during movement^24^, we reasoned that an enclosure with cues distributed along a single axis would exhibit the desired asymmetry. Below we characterise an anisotropy of such angular change in polarised environments.

Suppose we have a circular arena of radius *R* centred on the origin, with a polarising cue at distance *L* (Figure 1A). As an agent moves on a straight path b, we are interested in the angle *σ* from the agent’s heading to the cue, and how it changes when the agent moves. If the cue is within the arena (L<R), then the maximal change in angle (π radians) occurs when the agent moves towards and through the cue. If the cue is outside the arena (L>R), and the length of the path is very short (i.e. in the limit b → 0) we can calculate the change in angle *δσ* to the cue (see Figure 1A, middle panel for example illustration with a longer path):

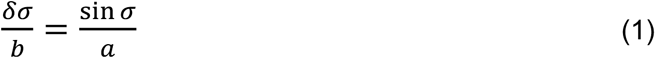

If the agent is on the x axis (i.e. *y* = 0) then we see that *δσ* is proportional to sin *σ* which is maximal for paths at 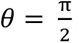, i.e. movements perpendicular (90°) to the cue.

Hence, spatial information during movement is not isotropic across directions (this conclusion holds on average for the entire arena). Specifically, angular change is maximal during movement perpendicular to polarising cues. Spatial computations, such as Euclidean triangulation, benefit from this parallax information and become more noise resilient (Figure 1 – figure supplement 4-6).

To corroborate this analysis, we conducted biologically inspired simulations of Euclidean triangulation in polarised environments (see Methods). As expected, the impact of spatial uncertainty was minimised for movement perpendicular to the polarisation axis (Figure 1 – figure supplement 4-6; two-sided Wilcoxon signed-rank test: Z= 1026.42, p<0.001). Results were robust within a plausible range of parameters.

Hence, in a circular environment polarised by two cues, the axis of greatest uncertainty corresponds to movement parallel to the axis defined by the cues. Thus, we would expect to observer larger errors in spatial memory responses in this direction than perpendicular to it. Conversely, to minimise positional errors, grid-patterns should orient to lie perpendicular to the polarisation axis.

### Effects of motion parallax on behavioural distance estimation

To test if participants can use anisotropic parallax information to improve distance estimates, we conducted a behavioural distance estimation task (N=20). In a sparse environment (Figure 2A, Figure 2 – figure supplement 1) polarised by two cues defining an axis, participants freely navigated to a start location. There they could initiate forward teleportation along one of three directions (−30°, 0° and +30° relative to the polarisation axis; Angles < 90° were chosen to allow testing of many distances with limited field-of-view, see Materials and Methods; Figure 2B; Figure 2 – figure supplement 1). At the goal location they gave an estimate of the traversed distance (Figure 2A, see Materials and Methods). As predicted, performance was most accurate when motion parallax was present. That is, when participants moved ±30° oblique to the polarisation axis, as opposed to along it. Figure 2C, absolute error, paired, two-sided t-test N=20, T (19) = 2.7, p=0.007; Figure 2 – figure supplement 2, mean error, T _(19)_ = 5.47, p<0.001).

**Figure 2.**
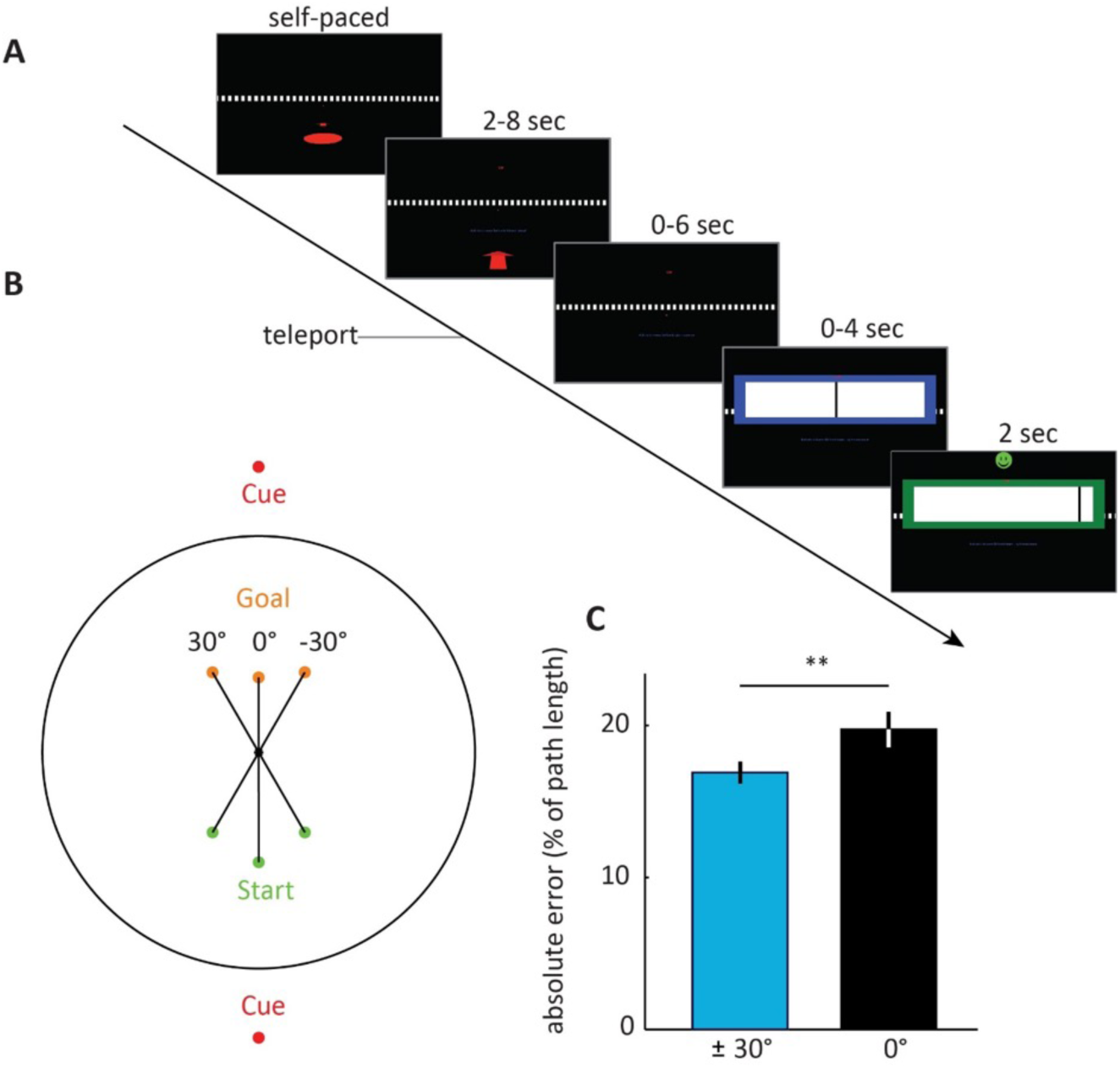
Behavioural experiment - distance estimation is most accurate on oblique paths. **A** Trial event sequence. Following a familiarisation phase, participants navigated to a start location (indicated by the red circle and arrow) and initiated teleportation in a given direction, either along a polarisation axis or at ±30° offset, see B and Figure 2– figure supplement 1. Teleportation distance was experimentally manipulated, and participants gave a distance estimate at the goal location by sliding a response bar (black slider in blue box). The cue was visible both at the start and the goal location (small red dot at eye height, in this example trial shown at the direction the arrow is pointing). Subsequently, participants received feedback. **B** Schematic of the three possible path angles shown at the same distance. Path distance varied from trial to trial (see Materials and Methods). Note that no boundary was present. The black circle only illustrates an analogy to the arena environments used in the fMRI experiments. Start and goal positions are illustrated by green and orange dots, respectively. **C** Distance estimation was most accurate on oblique paths, consistent with anisotropy of spatial information. Error bars show S.E.M. over participants.

### Polarised environments affect spatial navigation error

In order to test the effects of polarising cues in more naturalistic settings, we conducted navigation tasks in two distinct virtual environments (environment 1: N=50; environment 2 N=24). Both environments contained a grassy plain bounded by a cylindrical cliff, surrounded by polarising extra-maze cues (Figure 3). In environment 1, highly similar extra maze cues were visible in all direction (12 in total), but these changed orientation at two opposing points. These inversion points constituted a polarisation axis. A subset of participants that navigated in environment 1 underwent concurrent fMRI scanning (fMRI experiment 1; N=26), while another subset underwent concurrent eye tracking (eye tracking experiment; N=34). Environment 2 was more clearly polarised by a single cue on either side (see Figure 3 and Materials and Methods), with participants undergoing concurrent fMRI scanning (fMRI experiment 2; N=24). In each environment, participants performed a continuous object-location memory task^13, 19^ with 6 or 4 object-location associations (see Materials and Methods). Each trial comprised navigating to a target location - participants were able to move forward and rotate but not move backward - giving a response, and receiving feedback. This task was interrupted by occasional inter-trial-intervals when a fixation cross was presented on a grey screen for 2 seconds (on average after every third trial; range: 2-4). Object identity and location was randomised across participants (see Materials and Methods).

**Figure 3.**
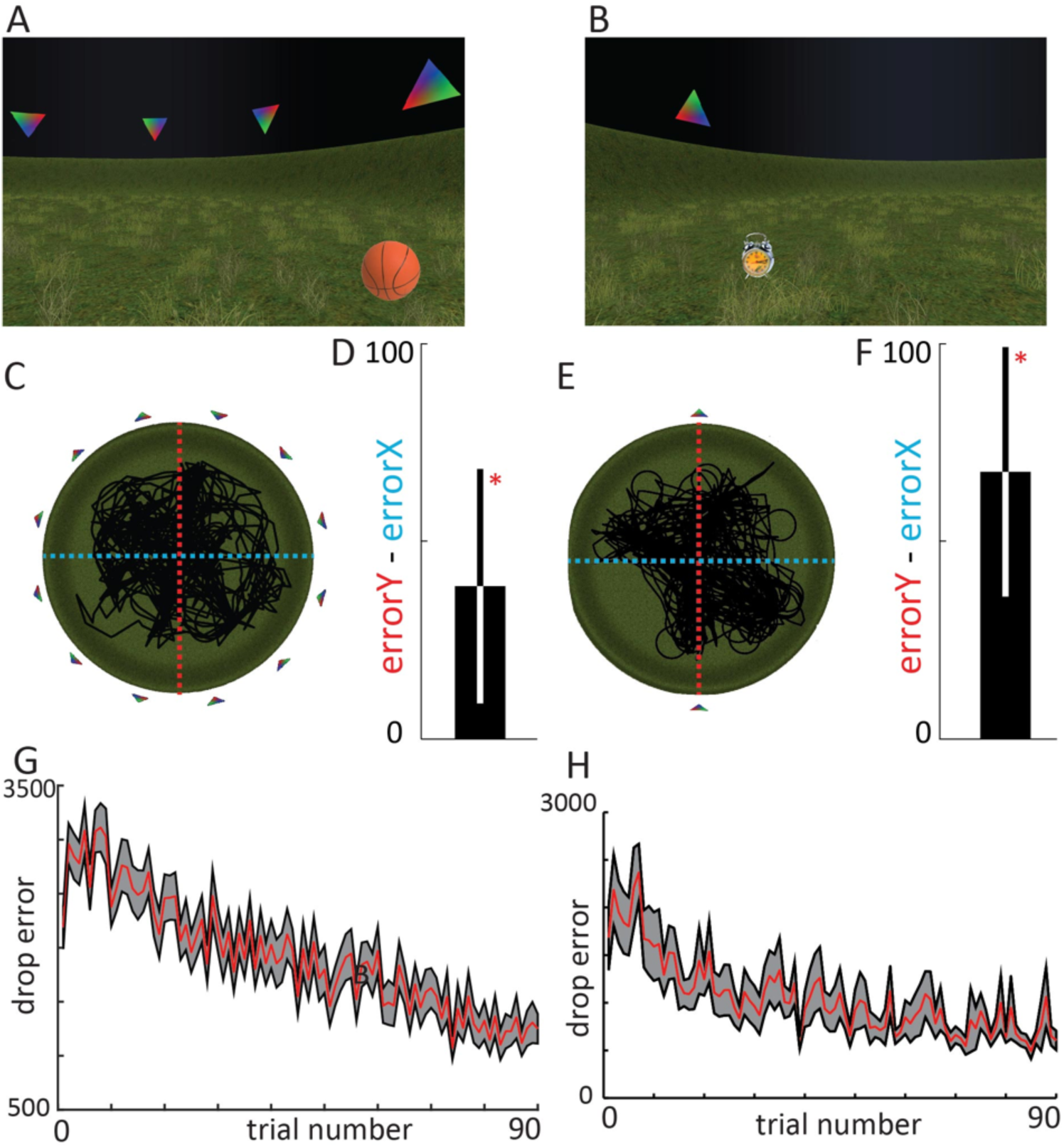
Spatial memory performance is anisotropic. Top. First-person view **A** environment 1 (used in fMRI experiment 1, and the eye-tracking experiment), **B** environment 2 (used in fMRI experiment 2). **C, E** aerial view. Human participants performed a free-navigation, object-location memory task (one example object shown on grassy plane, see Materials and Methods). In environment 1 an implicit polarisation axis was defined through the configuration of cues, i.e. the switch between upright and downward triangles. environment 2, an explicit polarisation axis was defined with two triangular cues alone. Black lines in aerial view show the paths of exemplary participants. Red dashed line indicates the polarisation axis (Y dimension), whereas the cyan dashed line indicates the orthogonal X dimension. **D, F** Bars show the median difference in spatial memory performance on the Y axis (i.e. the polarisation axis) versus the X axis. To avoid potential bias, we matched the number of trials in which participants faced (or moved) parallel and perpendicular to the polarisation axis (±45°). Spatial memory performance was anisotropic, with larger errors along the polarisation axis in both environments, which corroborates the theoretical predictions of an anisotropy in spatial information (Equation 1, Figure 1 and Figure 1 – figure supplement 4). **Bottom** Error bars show S.E.M. over participants.

Despite a relatively sparse environment, participants successfully learnt the object locations (Figure 3 – figure supplement 1). To avoid potential bias of the anisotropy measure due to the limitation of navigating using only three buttons, we matched the number of trials in which participants faced (or moved) parallel (y-axis) and perpendicular (x-axis) to the polarisation axis ±45° at the time of the spatial response (median difference in number of trials facing Y - number of trials facing X: environment 1 = 2; environment 2 = 16). We tested if participants’ spatial responses were more accurate when given perpendicular to the polarisation axes than in parallel with it, as predicted by the anisotropy in angular change information (Figure 3). To this end, for environment 1, we employed a linear mixed-effects regression model using the lmer function from the lme4 statistics package^25^ implemented in R 3.5.1 (R Core Team, 2018). As fixed effects, we tested the intercept of the effect of the median X error on the median Y error. As random effects, we used intercepts for the factor experiment with two levels (fMRI experiment 1 and eye tracking experiment; One-sided test; T _(57)_ =1.812, p=0.0376; Median percentage of X error relative to Y error = 3.8 %).

For environment 2, we employed a paired one-sided t-test (T _(23)_=2.1441: p=0.02141; Median percentage of X error relative to Y error = −15.2%). Participants’ spatial memory performance in both environments hence indicated that movement directions parallel to the polarisation axes were associated with low spatial certainty. Next, we asked if participants’ viewing behaviour would reflect increased exploration of more informative cues.

### Polarising cues are viewed for longer

We asked next if participant’s viewing behaviour would reflect the use of the polarisation axis. Notably, the visual appearance of all cues was matched and cannot explain any potential viewing time differences. We examined these potential differences in viewing times between cues using a repeated-measures ANOVA. Indeed, average percent viewing time differed between cues (Figure 4; F _(11)_ = 4.98, p<0.0001, n=34). However, our specific hypothesis was that the configural cues forming the polarisation axis (i.e. two pairs of cues of opposite orientation) were the ones most viewed and informative for navigation behaviour. A one-tailed paired t-test revealed that the configural landmarks were indeed viewed longer than the ones orthogonal to the polarisation axis (T _(33)_ =3.60, p=0.0005). This reliance on polarising cues correlated with spatial memory performance across participants. In particular, the difference in viewing time of polarising cues versus orthogonal cues correlated negatively with participant’s mean spatial memory error (Figure 4D. Pearson correlation, one sided, R= −0.2997, P = 0.0425).

**Figure 4.**
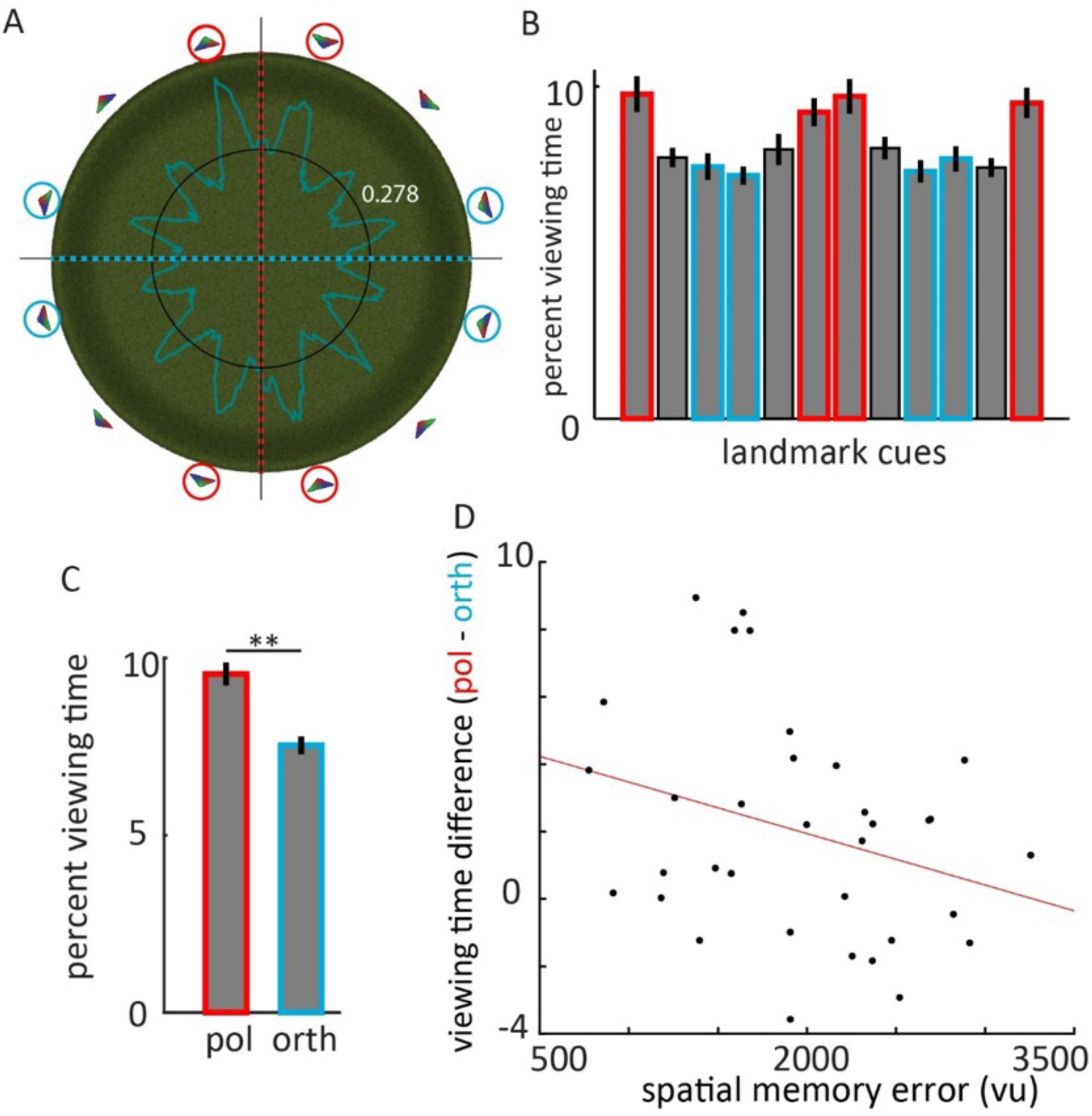
Eye tracking experiment -. Participant’s behaviour in the ‘12 cue’ environment (used in fMRI experiment 1 and the eye tracking experiment) reflects stronger reliance on the polarising cues. **A** Percent viewing time is plotted for fixations of 360 evenly spaced points on the boundary of the arena defined by the surface of the cues. Black circle: chance level of even distribution of viewing time across all points. Cues marked with red circles constitute the polarisation axis, cues marked with cyan are perpendicular to the polarisation axis. **B** Bars show average percent viewing time in 30° wide bins that were centered on each of the 12 cues. **C** Participants viewed the cues that form the polarisation axis longer than those perpendicular. **D** Spatial memory error correlated with the difference in viewing time of polarising cues versus orthogonal cues. Error bars show S.E.M. over participants.

The longer viewing times of the polarising cues and the correlation with spatial memory strongly suggests that they were key elements for. If hexadirectional activity as an index of grid-cell-like representations would exhibit a preferred orientation orthogonal to the polarisation axis, this would provide evidence for an adaptive nature of the impact of environmental geometry on the grid pattern.

### Grid-cell-like representation orient to misalign with an axis of high spatial uncertainty

To examine how grid-cell-like representations orient in simple polarised environments, we estimated hexadirectional entorhinal activity ^13, 15, 19^ for each participant. In brief, the method takes advantage of a six-fold periodic directional modulation of fMRI activity in entorhinal cortex during virtual movement (see Materials and Methods). First, we estimated individual orientations of hexadirectional entorhinal activity on the first half of the data. These orientations were not randomly distributed, but clustered approximately perpendicular to an axis defined by the configural cues (fMRI experiment 1, Figure 5). The absolute angle to the nearest ‘grid axis’ was approximately 30°, corresponding to maximum mis-alignment.

**Figure 5.**
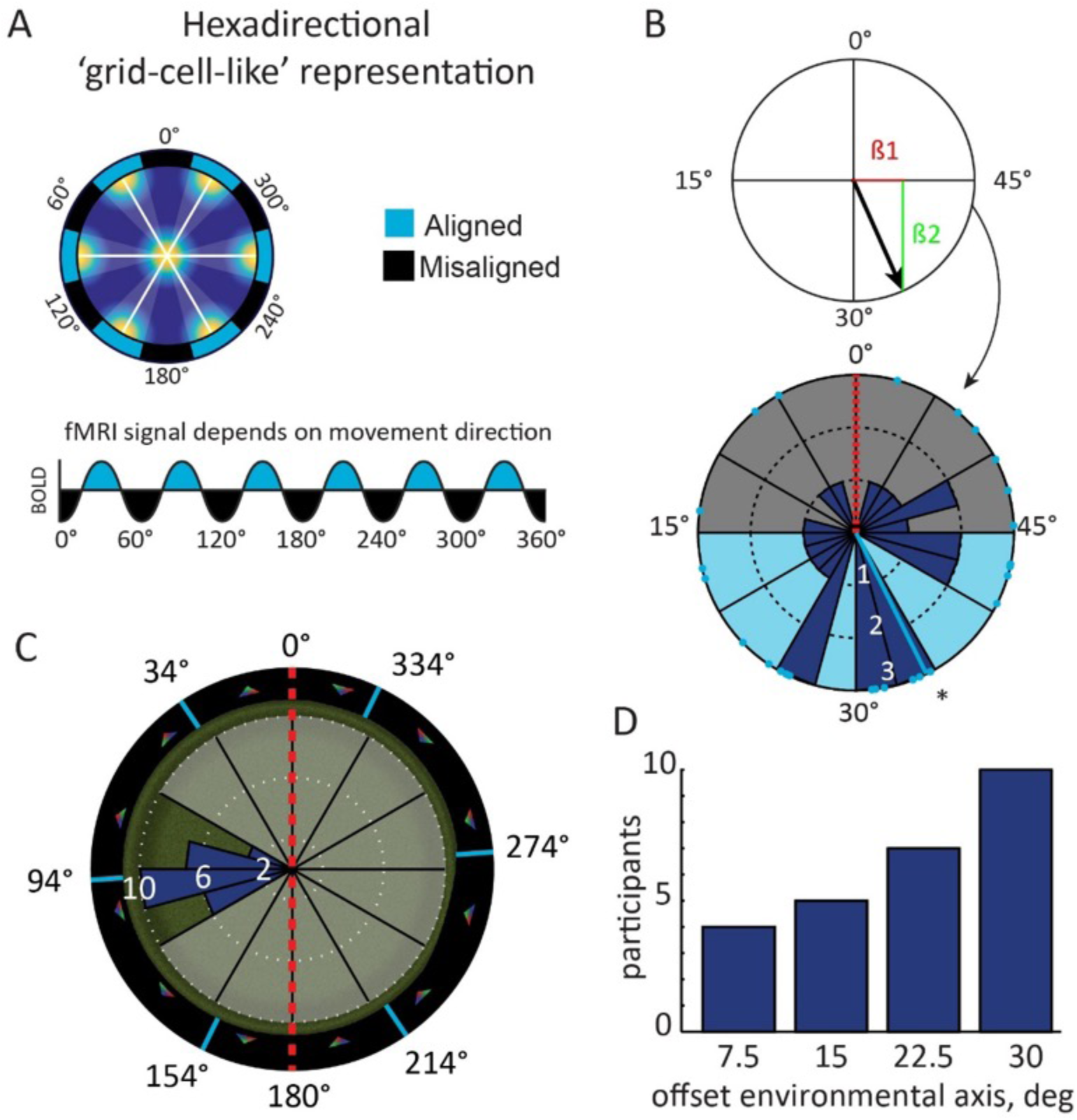
Hexadirectional activity in entorhinal cortex aligns perpendicular to the polarisation axis. **A** Hypothesis: movement parallel with the axes of grid cells is associated with increased fMRI BOLD signal ^16, 19^; **B** Top: Analysis procedure: the preferred orientation of hexadirectional fMRI activity in the entorhinal cortex was estimated by first fitting a general linear model (GLM) to the data with 60°-periodic sine and cosine regressors. This yields the associated parameter estimates ß1 and ß2, respectively. The preferred orientation in 60°-space (black arrow) can be derived from ß1 and ß2 (see Materials and Methods). The corresponding preferred orientation of hexadirectional activity in 360°-space can then be deduced. Here, this corresponds to multiples of 60° centered on 34° (light blue lines in C) relative to the polarisation axis (red dashed line) at 0°. **B** Bottom: Individual, preferred orientations in 60°-space (light blue dots) in right entorhinal cortex clustered at roughly 30° offset relative to the polarisation axis (red dashed line); mean orientation = 34° (light blue line). **C** Histogram of preferred hexadirectional activity plotted in full circular space (360°). Note that one of the hexadirectional axes (light blue lines) is roughly orthogonal to the polarisation axis (red dashed line), in line with optimal angles for self-localisation (Figure 1D) **D** Absolute angle between nearest axis of hexadirectional activity shown in B and the polarisation axis illustrate a tendency towards maximal misalignment. Note that the maximum offset is 30° due to the 60° periodicity of hexadirectional activity.

Circular mean = 34°; Figure 5C-D; N=26, circular V test for deviation from homogeneity perpendicular to the polarisation axis: V=6.68, p=0.032). Note that low-level visual features were equal in all viewing directions. Second, we performed a whole-brain analysis on the second half of the data. This confirmed that activity in right entorhinal cortex was increased for runs at periods of 60° aligned with the orientation identified from the first half of that data (Figure 6A-D, peak voxel t-test, T (25) =4.44, p=0.034, small-volume FWE-corrected). Consistent with the presence of grid-cell-like representations, runs aligned versus misaligned show largest activity increase for 6-fold rotational symmetry but not for biologically implausible control models of 5- or 7-fold rotational symmetry (repeated-measures ANOVA: F(3,25) = 8.3, p < 0.001; Post-hoc, paired t-tests with Holm-Bonferroni correction, * p<0.05). No other peaks remained across the cerebrum even at more liberal thresholds (p<0.001 uncorrected; T>3.45) and neither was there a notable circular clustering of hexadirectional activity in 2 control regions (mammillary bodies, which are close to the hippocampal formation: V test: V=3.33, p=0.822; right, primary visual cortex: V test: V=0.10, p=0.489; See Materials and Methods).

**Figure 6.**
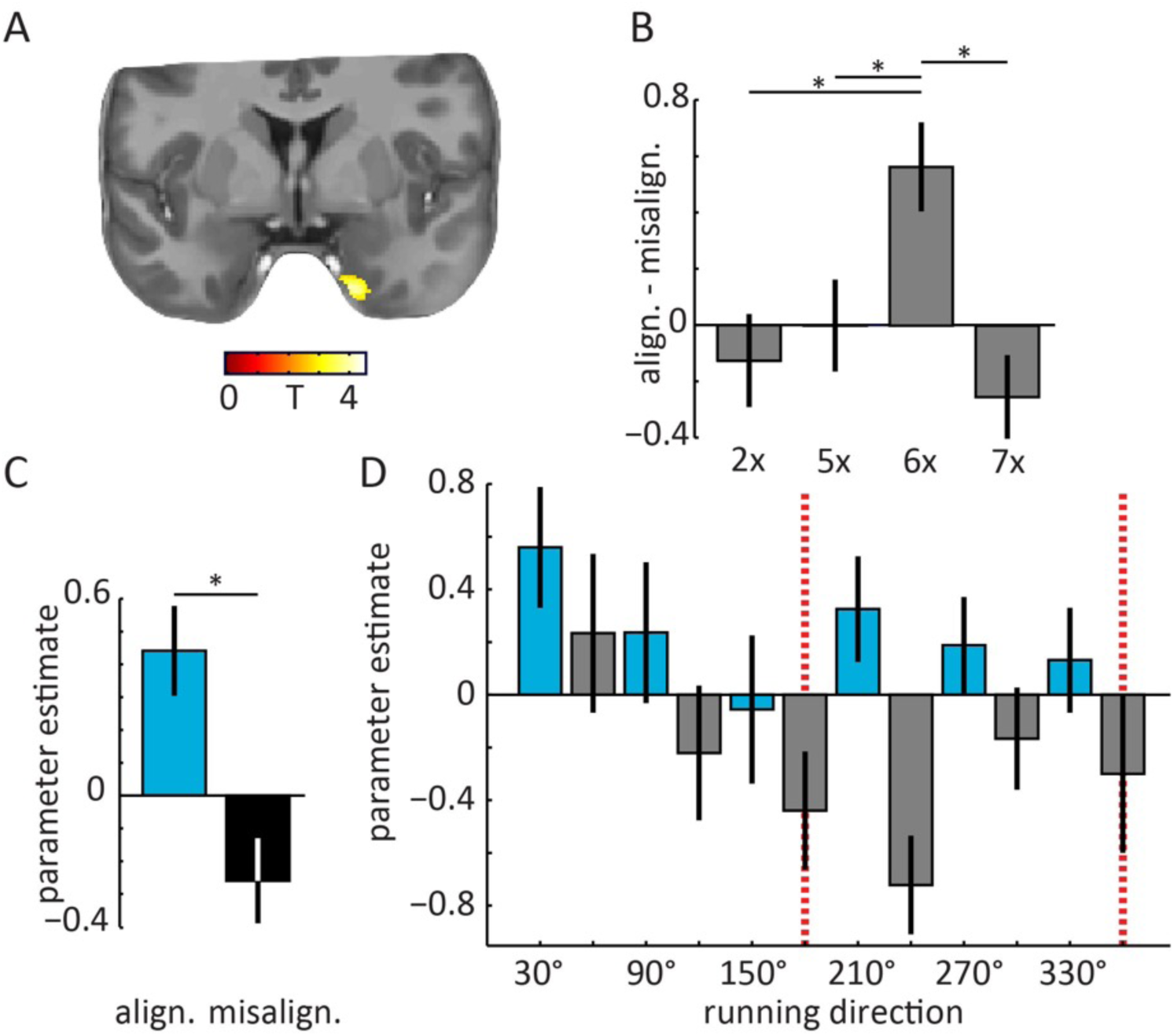
Cross-validation of hexadirectional activity in entorhinal cortex and control models. **A** A whole-brain cross-validation confirmed that entorhinal activity was increased on runs aligned with the predicted grid (i.e. runs in 30°-wide bins centred on 30°, 90°, 150° etc. indicated by light blue arrows in C and D; see Materials and Methods for details). Peak voxel t-test, T(25)=4.44, p=0.034, small-volume FWE-corrected). Image is thresholded at p<0.001 uncorrected for display purpose. Across the cerebrum no other peaks were observed at this threshold. The T statistic (colour bar) is overlaid on the structural template. **B** In agreement with grid-cell-like representations, runs aligned versus misaligned show largest activity increase for 6-fold (6x) rotational symmetry but not for biologically implausible control models. Next to 5- and 7-fold rotational symmetry, 2-fold symmetry was tested to rule out a direct effect of running parallel to the polarisation axis or not. For all analyses the aligned condition was centered at an angle equivalent to 90° from the polarisation axis (e.g. 30° for 6-fold symmetry). **C** Parameter estimates of runs aligned (light blue, see schematic grid in Figure 1C right panel) and misaligned (black) with the predicted hexadirectional orientation extracted from the peak voxel in A. **D** To examine the influence of different running directions, we plotted the parameter estimates for separate regressors of 12 directional across the entire time-series of fMRI data from the peak-voxel in A. Note the alternating pattern of activity aligned and misaligned. Red dashed lines indicates the polarisation axis. Bars show means and S. E. M. across participants.

To test if the environmental anchoring depends on the configural cues, we scanned another group of participants in an environment with a non-configural, polarisation axis consisting of only two extra-maze cues (fMRI experiment 2). We estimated individual orientations of hexadirectional activity, and again found that they clustered perpendicular to the polarisation axis (circular mean = 32.28°; Figure 7) replicating the findings from the first experiment (N=24, circular V test: V = 5.95, p=0.043; Figure 7). Sampling of running directions could not explain these effects in either experiment (Figure 3 – figure supplement 1). In sum, the results from fMRI experiment 1 and the replication in fMRI experiment 2 provide converging evidence that the preferred orientation of hexadirectional activity in entorhinal cortex depends on navigation-relevant, polarising cues, independent of the specific type of cue (configural or non-configural). The orthogonal arrangement of hexadirectional activity, as an index of grid-cell-like representations, is in agreement with optimal activity patterns of simulated grid cells for self-localisation, suggesting that the impact of environmental geometry on grid cells may be adaptive.

**Figure 7.**
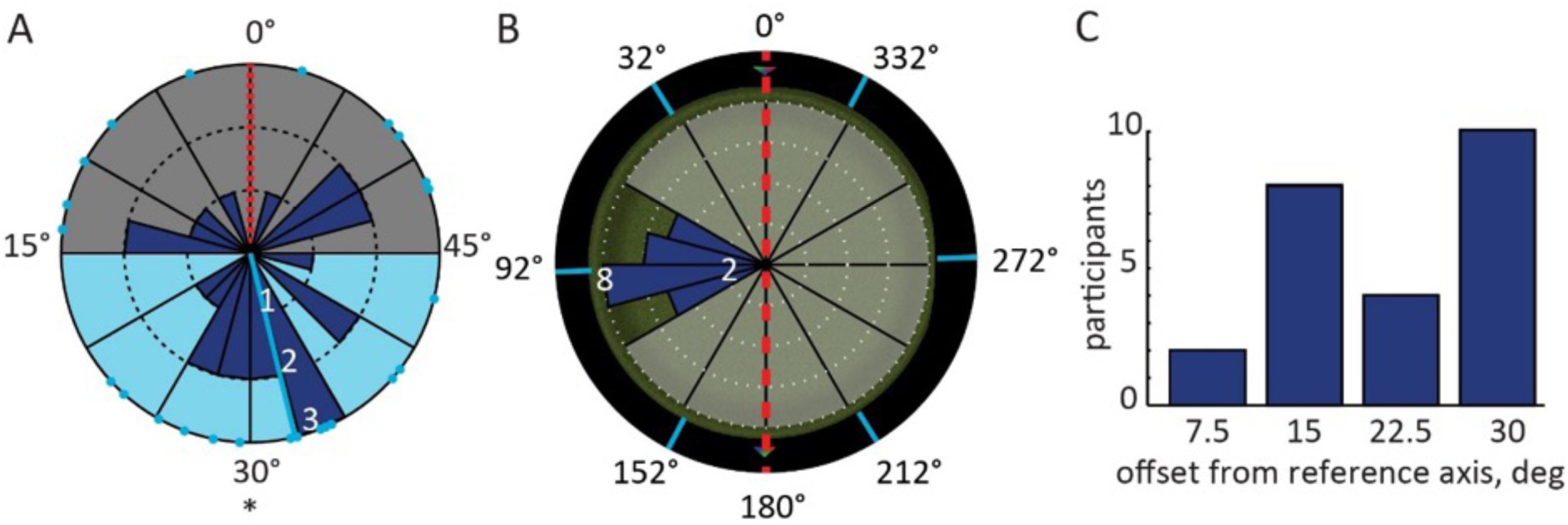
Environmental effects on hexadirectional activity in fMRI experiment 2. **A** Preferred hexadirectional activity in 60°-space (light blue dots) in right entorhinal cortex clustered at roughly 30° offset relative to the polarisation axis (mean orientation = 32.28°; light blue line), in line with optimal angles for self-localisation (Figure 1D) **B** Histogram of preferred hexadirectional orientations plotted in full circular space (360°) onto a top-down view of the arena. Note that one of the putative grid axes (light blue lines) is roughly orthogonal to the polarisation axis (red dashed line). **C** Absolute angle between nearest axis of hexadirectional activity shown in B and the polarisation axis. Note that the maximum offset is 30° due to the 60° periodicity of hexadirectional activity.

## DISCUSSION

We examined the effects of polarising cues on spatial navigation behaviour and grid-cell-like representations. We found that estimation of movement distance in a virtual environment was least accurate when participants moved along a polarisation axis and no motion parallax was present. Likewise, spatial memory performance in free-navigation, object-location memory tasks in polarised environments showed relatively larger errors parallel to the polarisation axes (i.e. anisotropy). Eye-tracking recordings indicated that participants viewed polarising cues for longer than other cues, which correlated with their spatial memory performance.

To test the theoretical implications of anisotropic spatial information on a system of grid cells, we used biologically inspired simulations. We demonstrate that the representation of self-location is most accurate when grid-patterns align with the axis of highest spatial information. Motion induced parallax is a source of navigation-relevant information^24, 26^ and lies at the heart of surveying unknown terrain for the creation of spatial maps. With respect to parallax, the optimal grid-pattern alignment corresponds to grid angles 30° offset from the polarisation axis. fMRI-based estimates of grid-cell-like representations^13, 14, 16–19^ showed consistent orientations across participants in two independent experiments, yielding the optimal grid orientation for decoding self-location. Taken together, our results provide evidence that the grid system aligns to axes of high spatial information, which suggests that effects of environmental geometry on the grid system are adaptive responses in service of flexible navigation. Furthermore, angular change to stationary cues during self-motion may play a central role in the computations underlying the grid system.

Could grid-cell-like representations simply follow extra maze cues or object locations? Note that the peaks of hexadirectional activity (grid-cell-like representations) did not correspond to movement directions directly facing a cue in either fMRI experiment. This speaks against a simple sensory ‘anchoring’ of grid-cell-like representations to landmarks in the environment. Importantly, the coherent orientations of grid-cell-like representations can also not be attributed to the presence of objects in each trial, since their locations were randomised across participants and the two fMRI experiments had different numbers of objects.

Our findings are in agreement with reports showing that entorhinal fMRI activity correlates with Euclidean distance to a goal location ^27^ and the proposal that grid cells might enable goal-directed vector navigation ^1, 2, 5, 28–32^. Interestingly, both angular and distance information that is needed for triangulation can be derived from either visual or proprioceptive and vestibular cues. For example, visually modulated cells in the rat posterior-parietal cortex signal the egocentric cue direction ^33^ and head-direction cells in the entorhinal cortex and other regions realign to visible cues, but also function without vision and rely on vestibular information ^34^. On the other hand, distance information can be inferred visually from the relative size of objects and cues ^24^ or is based on proprioceptive and timing information during movement, both of which modulate grid cell activity ^35^. Hence, triangulation for navigation could bridge different sensory modalities. Furthermore, it combines egocentric cue directions and distance information to infer map-like, survey representations of the environment, thereby naturally integrating egocentric and allocentric reference frames, which are not mutually exclusive and can work in parallel and across brain regions ^33, 36, 37^.

A potential avenue for future studies is to examine the effect of anisotropic spatial uncertainty on rodent grid-cell firing, albeit many animals may be needed to detect such a stochastic effect. It is known that grid cell firing exhibits plasticity, regularising and reorienting incrementally with continued experience of an enclosure (Barry et al., 2007; Barry, Heys, & Hasselmo, 2012; Carpenter et al., 2015; Stensola et al., 2015). Our results suggest that these changes likely optimise the grid-code, allowing for an increasingly accurate representation of self-location. However, the physiological and circuit mechanisms that facilitate and direct this process are currently unknown. Theoretically, a number of authors have considered the impact of noise in grid cell coding of self-location and its implications for the capacity and error-tolerance of the entorhinal spatial representation ^5, 41, 42^. However, to the best of our knowledge, we have provided the first theoretical and practical account of anisotropic spatial uncertainty on the grid system. It remains to be seen if such asymmetries, which are likely a common feature of the environment ^43^, exert more wide-ranging influences on grid-firing; distorting the grid-pattern or changing the relative scales of different grid modules; or might also impact grid-like coding of non-spatial information. In conclusion, we combined biologically inspired computational models, behavioural tests, eye-tracking and fMRI-based proxy measures of grid-cell-like population activity to test the effects of environmental geometry on the entorhinal grid system. Our results are consistent with an adaptive and flexible role of grid cells in self-localisation and navigation. This opens up the exciting possibility for a deeper understanding of fundamental neural building blocks of cognition and behaviour.

## Materials and Methods

### Simulation of Euclidean triangulation

To test the impact of stochastic fluctuations or noise on triangulation accuracy, we implemented the following simulation in Matlab (2012b, The MathWorks Inc., Massachusetts). Triangles were formed by two points representing start and end points of a straight path in the horizontal plane (e.g. observer locations at time point 0 and time point 1) and one of two polarising, stationary cues. Triangulation was based on the sine rule according to:

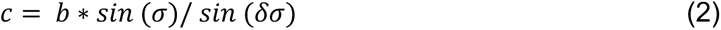

where *c* is the unknown side (distance to the cue at the end point; Figure 1 – figure supplement 4), *b* the known side (distance travelled), *σ* the angle to the cue at the start point and *δσ* the angular change to the cue between start and end point.

Path orientation (azimuth) was varied in steps of 1°, path length remained constant and each path was centred on the origin of the coordinate system. Hence, the start and endpoints of different paths mapped onto a circle. This ensured that the mean distance of different paths to one or multiple cues remained constant. Before the triangulation iterations, random noise was added to the known side and the two distance angles. The error in side length had a mean absolute deviation of roughly 5% the original length (based on typical human distance errors during walking ^44^) and was drawn from a Gaussian distribution with mean 0 and a sigma of path length of 15.95. The absolute angular error for a single angle was 5° on average (drawn from a von Mises distribution with mean 0 and a sigma of 6.26) and 15° on average for the absolute cumulative error across all three angles of a triangle. This error rate was based on the mean, absolute angular error observed in humans performing a triangle completion task in virtual reality, which involved pointing to a start location after an outward path with two turns ^45^.

#### Triangulation measurements: noise resilience

Triangulation was repeated for all sides of a triangle using the known base *a*. If the inferred side was the base (the path), triangulation was repeated with both remaining sides serving as the known side and the two results were then averaged. Dual triangulation for the base was done to avoid biased results due to the selection of any one of the remaining sides. Note that the length of the remaining sides was not constant and changed in opposite directions for different path angles, potentially affecting the noise resilience measure at different path angles. This was not a problem in the reverse case, because the base (side *a*) had constant length. The triangulation error for the 3 sides was computed as the absolute difference in the original side length and the length based on triangulation with noisy input parameters. The 3 error rates were then averaged for further computations and the assessment of noise resilience across paths (Figure 1 – figure supplement 4-6). Furthermore, the distance between the most proximal cue to the centre of each path (the middle of the base of a triangle) was always equal to the length of the path, with the exception of Figure 1 – figure supplement 6 that shows the effects of different path lengths and different noise levels. In other words, usually the path length was half the length of the polarisation axis. Triangulation to additional cues was performed for a given path angle if these were within ± 90° (determined from the centre of a path) to emulate a limited field-of-view. This meant that cues in only that half of the environment were used for triangulation that was faced on a given path (1 point in Figure 1 – figure supplement 4).

#### Triangulation measurements: triangle quality

The quality measure for triangle shape (triangle area divided by the sum of squares of side lengths; Figure 1 – figure supplement 4 light blue curve) was modified from ^46^ who describe optimization of finite element triangulations in the generation of meshes.

### Computational models of grid cell systems

#### Grid cell system model

Spiking activity of a population of grid cells, organised into 4 (except where otherwise specified) discrete modules by spatial period size, was modelled in a two-dimensional circular environment of radius 50cm using Matlab v.8 (Mathworks). Spatial periods or grid scales, *λ_i_*, were determined as a geometric sequence beginning with *λ_1_* = 25cm and increasing with a scale factor of 1.4 (except where otherwise specified). Tuning curves for each grid scale *λ_i_* were generated with locations of grid nodes specified as a regular triangular grid and expected firing rate at each location determined by a Gaussian distribution centred on the nearest node:

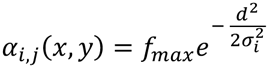

where *j* specifies a particular cell, *d* is the distance from (x,y) to the nearest grid node, *f_max_* the maximum firing rate (constant across the population; 10Hz except where otherwise specified), *σ_i_* the tuning width of the grid fields 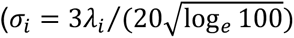 following^4^.

Within each of the 4 modules, M = 195 offset tuning curves were distributed in a 13 × 15 rectangular grid via translations of this original tuning curve, as well as adding a random translation common to all grids in the module. This resulted in a total of 1560 grid cells in a system. Grid tuning curves could also be rotated to specified orientations; all grid tuning curves always shared a common orientation. All these transformations were performed using cubic interpolation.

In each iteration of the model, the true position (x, y) was specified as the centre of the circular environment (0, 0). To model uncertainty, Gaussian noise, with standard deviation varied independently in x and y, was generated separately for each module and added to (x, y), to yield a noisy position estimate (x + ε_x,i_, y+ ε_y,i_). Anisotropic uncertainty was produced by independently varying the standard deviations of ε_x,i_ and ε_y,i_ between 0 and 5. All cells within a module therefore received the same noisy position input, but cells in different modules received different input. Thus cell firing rate was now modulated according to α_i_ (x + ε_x,i_, y+ ε_y,i_).

Higher-order “four-leaf” and “six-leaf” noise distributions were created for control experiments (Figure 1 – figure supplement 3). In polar coordinates, the width (s.d.) of the Gaussian noise distribution along the ray was modulated by a cosine function of the angular coordinate.

The signal extracted from the grid cell system was the number of spikes, *k*, generated by each neuron during a finite read-out period, *T* = 0.1s (the approximate length of a theta cycle) – i.e. a population response ***K*** *= (k_1_, …, k_N_).* We assume the decoding cannot take the added noise into account in any way, so that given a position *x* the probability of observing the response ***K*** in time T, following ^4^, is taken to be:

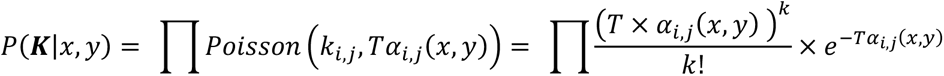

where *α_i,j_(x,y)* is calculated by cubic interpolation from the tuning curve. From the population response ***K***, we can decode position as the maximum likelihood estimate of *(x, y)*, that is *x̂,ŷ(**K**).* Given the initial assumption that all positions within the environment are uniformly likely,

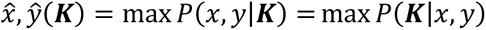

Thus *x̂,ŷ(**K**)* may be closely approximated by calculating *P(**K**|x,y)* for a sufficiently finely spaced uniform sample of *x* and *y* values across the environment, and selecting the values of x and y which yield the greatest *P(**K**|x,y)*. We used a spatial bin size of 0.5 cm. Where two or more solutions yielded the same maximal *P(**K**|x,y)* (i.e. decoding was ambiguous), one was randomly selected ^2, 4^.

#### Assessing grid system performance

For each combination of levels of uncertainty in x and in y, we assessed the performance of grid systems whose patterns were orientations to these x-y axes from 0° to 30° at intervals of 2.5°. For each case, five experiments each consisting of 15,000 iterations of this procedure were performed. In each of these five experiments, the square grid across which the environment was sampled to produce tuning curves was set at a different orientation to the environment’s Cartesian axes, in order to control for any effect of uneven sampling (the orientations were 0° and 4 orientations randomly selected and then used across all conditions). The results of equivalent pairs of uncertainty levels (e.g. standard deviation respectively in x and y of 0 and 5 cm, and 5 and 0 cm) were combined to total 2 × 5 × 75,000 = 150,000 iterations. Using these, accuracy of decoding was assessed via the approximated maximum-likelihood estimate square error, or MMLE, based on the square errors of position decoding:

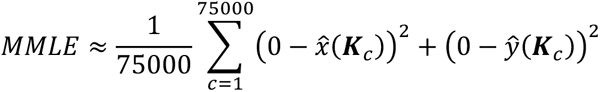

### Neuroimaging, behavioural- and eye-tracking experiments

#### Participants

##### FMRI experiment 1

26 participants took part in the study (12 females, age range: 19–36, mean age: 23 years). Materials and Methods were approved by the local research ethics committee (ethics committee University Duisburg-Essen, Germany and CMO region Arnhem-Nijmegen, NL). Written informed consent was obtained from each participant.

##### FMRI experiment 2

25 participants took part in this study (11 females, age range: 18-32, mean age: 24 years). One participant was excluded from the analysis due to poor performance (i.e. 55 trials with no location response within 30 seconds of the respective trial, more than a standard deviation above the mean). Materials and Methods were approved by the local research ethics committee (CMO region Arnhem-Nijmegen, NL). Written informed consent was obtained from each participant.

##### Behavioural experiment

20 participants (11 females, age range: 18-24, mean age: 20 years) participated in the behavioural experiment. Materials and Methods were approved by the local research ethics committee (CMO region Arnhem-Nijmegen, NL). Written informed consent was obtained from each participant.

##### Eye tracking experiment

36 participants (15 females, age range: 18-63, mean age: 26) participated in the experiment. Two participants aborted the experiment early, because of VR-induced nausea, and their data was excluded from all analyses. Materials and Methods were approved by the local research ethics committee (CMO region Arnhem-Nijmegen, NL). Written informed consent was obtained from each participant.

#### FMRI acquisition

##### FMRI experiment 1

Blood-oxygenation-level-dependent (BOLD) T2*-weighted functional images were acquired on a 7T Siemens MAGNETOM scanner (Siemens Healthcare, Erlangen, Germany) using a three dimensional echo-planar imaging (3D EPI) pulse sequence ^47^ with a 32-channel surface coil with the following parameters: TR = 2.7 s, TE = 20 ms, flip angle = 14°, voxel size 0.9 × 0.9 × 0.9 mm, field of view (FoV) = 210 mm in each direction, 96 slices, phase encoding acceleration factor = 4, 3D acceleration factor = 2. The scanning session was subdivided into EPI acquisition blocks of 210 volumes each. The majority of participants performed 5 blocks over the course of approximately 55 minutes.

Deviations from the 5 blocks in a few participants were due to technical problems or interruptions on behalf of the participants (3 participants had 4 blocks, 2 participants 6 blocks). In addition, T1-weighted structural images (MP2RAGE; voxel size: 0.63 mm isotropic) and a field map (gradient echo; voxel size: 1.8 x 1.8 x 2.2 mm^3^) were acquired. Results of an entirely unrelated, task-independent whole-brain connectivity analysis of data from experiment 1 have been described in a previous report ^48^.

##### FMRI experiment 2

BOLD T2*-weighted functional images were acquired on a 3T Siemens Trio scanner (Siemens Healthcare, Erlangen, Germany) using a three dimensional echo-planar imaging (3D EPI) pulse sequence ^47^ with a 32-channel surface coil with the following parameters: TR = 1.8 s, TE = 25 ms, flip angle = 15°, voxel size 2 × 2 × 2 mm, field of view (FoV) = 224 mm in each direction, 64 slices, phase encoding acceleration factor = 2, 3D acceleration factor = 2. Each scanning session consisted of an EPI acquisition block of 1031 volumes on average (range: 661-1200), amounting to roughly 31 minutes of scan time. In addition, T1-weighted structural images (MPRAGE; voxel size, 1 mm isotropic; TR, 2.3 s) and a field map (gradient echo; voxel size, 3.5 x 3.5 x 2 mm^3^) were acquired.

#### Experimental tasks

##### FMRI experiment 1

Participants freely navigated a 3D virtual reality environment with a modified version of the arena from the studies by Doeller and colleagues ^13, 19^ (Figure 3A) using a 4-button controller. UnrealEngine2 Runtime software (Epic Games) was used to generate the virtual reality task. Instead of two orthogonal axes that are formed by the walls of square enclosures (as in (Krupic et al., 2015; Stensola et al., 2015)) we opted for the simplest case of a single axis, which was determined by extra-maze cues in a circular arena. We hypothesized that the orientation of grid representations would be coherent across participants, as shown in rats moving through square environments, and that this orientation would be determined by the amount of spatial information obtained on movement paths of such orientation. The environment consisted of a circular arena with 12 extra-maze cues, 6 upright and 6 inverted triangles. Two pairs of neighbouring triangles of different orientation comprised the two configural cues on opposite sides of the arena that defined a polarisation axis. To control for possible visual effects on our direction-related analysis, we designed the colour textures for the extra-maze cues in such a way, that the low-level visual features remained equal across cues. Each triangle had a red, green and blue corner, arranged in 1 of 6 possible constellations. The arrangement of textures was randomised across participants. Counting one’s steps was not possible, because no body parts were visible, and the virtual pitch direction was fixed. Self-motion information such as optic flow induced by the grassy plane was present but did not yield directional information. Participants performed a self-paced object-location memory task that involved collecting and replacing six everyday objects to locations that were randomised across participants. Participants collected each object from its associated location once during an initial phase, by running over it. Navigation was not interrupted during the transitions between trials to enable more natural (ecologically valid) continuous navigation. In each subsequent trial they saw an image (cue) of one of the objects in the upper part of the screen and had to move to the object’s associated location and press a button (replace phase). After this response, the object appeared in its associated position and participants collected it again (feedback phase). After an average of 3 trials (range 2-4 trials), a fixation cross on a grey background was presented for 4 seconds (inter-trial-interval, ITI). Object locations were randomised across participants. Since the task was self-paced, the number of trials varied across participants (range: 94-253; mean: 179). Prior to the fMRI experiment, participants performed a similar object-location task with different objects in a different virtual environment outside the scanner to familiarise themselves with the task demands.

##### FMRI experiment 2

Participants freely navigated the same virtual environment as used in fMRI experiment 1, but with only two extra-maze cues on opposite sides of the arena that defined a polarisation axis (Figure 3A). Participants performed the same object-location memory task described above, except that 4 objects were used instead of 6. Participants performed an average of 117 trials (range: 63-179). Prior to the fMRI experiment, participants performed a similar object-location task with different objects and a different virtual environment outside the scanner to familiarise themselves with the task demands.

##### Behavioural experiment

Participants freely navigated a virtual reality environment (Figure 2A, Figure 2 – figure supplement 1) by using four buttons on a keyboard to move in the four cardinal directions and the mouse to change horizontal viewing direction. The virtual environment was displayed at 1680×1050 pixel resolution and 60 Hz refresh rate approximately 40cm in front of the participants’ eyes. They were teleported between varying start and end locations at one of three possible angles and performed a distance estimation task. The environment was a ‘pitch black’ space with otherwise only three distinguishable elements. First, it included a background consisting of a white dashed line oriented horizontally and projected at infinity. This background provided minimal visual information to perceive rotational movements as well as motion parallax of a cue viewed from different angles. Second, a cue, consisting of a red circle, was displayed vertically on a fixed location. Third, a red circle indicated the start location of each path with an arrow pointing in the direction of the goal location. The rationale behind using a visually sparse environment and teleportation to the goal location was to prevent the use of other distance cues, such as cue size (e.g. patches of grass or a boundary) or an estimate of ‘time-of-flight’, respectively. This ensured that the change in size of the cue and the change in angle and motion parallax to the cue from start to the end of a path was the sole means by which the distance estimation task could be performed correctly. Prior to the experiment, participants performed a similar distance estimation task in a different virtual environment to familiarise themselves with the task demands. At the beginning of the behavioural experiment, participants were instructed to approach the cue in order to familiarise themselves with its location and distance.

The trial structure was as follows: Participants were instructed to navigate to the starting point. Once they reached the starting point, their movement was restricted to rotations and the message ‘click right mouse button to teleport ahead’ was displayed (orientation phase one). Participants could self-initiate teleportation to the goal location by a mouse-click and orienting towards the pointing direction of the arrow, at which point the view was frozen and teleportation commenced 2 seconds later. After teleportation to the goal location, the start location became invisible (the red circle with arrow disappeared), movement remained restricted and only rotations were possible and the message ‘click right mouse button to give response’ was displayed (orientation phase two). Participants could self-initiate the response phase. Then, a horizontally oriented window was displayed together with the message ‘indicate distance (left = minimum, right = maximum)’ and participants could move the mouse to slide a bar inside the window to indicate how far they thought they were being teleported.

The range of possible responses was 0 virtual units (vu) to 6000 vu. For comparison, the arena diameter used in the fMRI studies was 9500 vu for the inner boundary and the length of the polarisation axis (i.e. the distance between opposing, extra-maze cues) was 12064 vu. The range of teleportation distances was 500 vu to 5500 vu (mean = 2742 vu). The response was finalised by another mouse click and subsequently, feedback in the form of smiley faces was given for 2 seconds. The color of a smiley for a response error < 2% of the correct distance was green, light green for an error < 4 %, yellow for an error < 8 %, orange for an error < 16% and red otherwise. During this feedback phase, participants could still move the response bar to see other response-to-feedback mappings (i.e. the smiley associated with a given horizontal pixel location). Once the feedback disappeared, participants were able to freely navigate again. At the beginning of about 50% of trials (determined pseudo-randomly), participants were placed to a point in front of the start location to speed up the experimental procedure (i.e. to reduce navigation time from a goal location to the start location of the subsequent trial) and thereby increase the number of trials. In addition, the orientation phase 1 and 2 were restricted to 6 seconds and the response phase to 4 seconds indicated through the display of a timer. If the time limit was reached, ‘Time is up! This trial is invalid’ was displayed on a red background and no response was recorded. Teleportation distances and teleportation directions were pseudo-randomly determined on each trial. Teleportation directions were either 0° (approaching the cue on a straight line), −30° or +30°. The location of the cue was at (x = 0 vu, y = 8500 vu) and following the approach of the simulations, all paths were centered on the origin of the coordinate system. However, this would provide a relative advantage to the parallel condition. The size of the cue directly reflects its distance to the observer, which becomes particularly apparent at close proximity. In the −30° and the +30° conditions, the goal location is always further away from the cue compared to the 0° condition at equal teleportation distances. Furthermore, the independent measure (teleportation distance) is linearly associated with goal-to-cue distance only in the 0° condition. To avoid bias due to unequal goal-to-cue distance, we equalized this measure by subtracting the difference across conditions (at equal teleportation distances). In effect, this shifted the teleportation paths in the 0° condition backwards by a given amount (Figure 2B). Due to a limited field-of-view of 85°, testing of large path offsets of e.g. 90° was not feasible. The task duration was limited to 30 minutes in which participants performed an average of 129 self-paced trials (range: 52-238). Prior to the main task, participants performed a training version of the task in a richer virtual environment with a comparable trial structure where the length of the path was not traversed by teleportation but rather through guided movement.

##### Eye tracking experiment

During a magnetoencephalography study (MEG; data are subject of an independent report), participants performed the same task in the same virtual environment as in fMRI experiment 1 (i.e. the environment with 12 cues). However, they had to learn the locations of 8, instead of 6 objects. Simultaneously, gaze position and pupil area of the left eye were monitored with an infrared-based Eylink 1000 eye tracking system at 1200 hz.

#### Analysis of eye tracking data

The eye tracking data were converted to screen coordinates. Blinks were removed from the time series based on deviations in pupil area of more than one standard deviation from the mean including 25ms around the blink on- and offsets. After smoothing with a running average kernel of 10 ms and linearly detrending, gaze positions were transformed to velocities expressed as degree visual angle per second. Since gaze velocity profiles differed between translations and rotations during navigation, saccades were detected individually during head rotations and static or translational navigation. During head rotations, saccades were detected using a threshold of 12 times the median-based standard deviation of velocity, during static or translational periods with 6 times the median-based standard deviation of velocity. Saccades shorter than 12 ms were excluded ^49^.

To examine which cues were looked at most during the experiment, we transformed horizontal gaze positions to degrees visual angle and scaled them linearly to match the physical field of view (36 degrees visual angle) to the virtual field of view (85 degrees virtual visual angle). The resulting virtual degrees visual angles were then combined with the virtual head direction to reconstruct the allocentric viewing direction at each point in time. Since all twelve spatial cues were visible exclusively in the upper visual field, we limited our analysis to gaze positions on the upper half of the screen. Moreover, to account for any potential influence of initial viewing angle on this analysis, we excluded all samples recorded before participants rotated at least 90 virtual degrees away from the starting viewing angle (average time excluded at start of experiment: 15.2 seconds). We then computed the intersection between allocentric gaze and a surface at the radius of the extra-maze cues from the centre of the arena for each point in time as follows. We first generated 360 equally spaced vertices at the radius of the extra-maze cues and computing the vectors between the participant’s location and all of these vertices. We then selected the respective vertex with the minimal angular distance to the current allocentric viewing direction. This way, we obtained the position of gaze on the arena border or cues at each given point in time. For each vertex we computed the respective viewing time expressed as percent of all samples obtained for each respective participant to account for differences in experiment duration across participants (Figure 3 – figure supplement 2). To examine whether there were differences in viewing times between cues, we then binned vertices into 30-degree bins centred on the cues and compared average viewing time using a repeated-measures ANOVA. However, our specific hypothesis was that especially the cues forming the polarisation axis should be most informative for the task. To test whether these cues were the ones most viewed, we averaged viewing times for the configural landmarks that comprised the polarisation axis (i.e. two pairs of cues of opposite orientation) and compared it to the average of the four cues orthogonal to the polarisation axis using a one-tailed paired t-test.

### FMRI data pre-processing

Image pre-processing and analysis were performed with the Automatic Analysis Toolbox (https://github.com/rhodricusack/automaticanalysis). This involved using custom scripts combined with core functions from FSL 5.0.4 (http://fsl.fmrib.ox.ac.uk/fsl/) and SPM8 (http://www.fil.ion.ucl.ac.uk/spm). SPM was used for an iterative functional image realignment and unwarping procedure to estimate movement parameters (three for rotation, three for translation) and to correct images with respect to gradient-field inhomogeneities caused by motion. To improve co-registration and the creation of a group-specific structural and functional template using the Advanced Neuroimaging Toolbox (ANTS; http://www.picsl.upenn.edu/ANTS/) structural images were de-noised using an optimised non-local means filter ^50^ and mean EPI images were corrected for gradual changes in signal intensity (bias correction) using SPM. Next, structural images were co-registered (based on mutual information) to the functional images using SPM and brain-extraction was performed using FSL. The resulting skull-stripped structural image was segmented into grey matter (GM), white matter (WM) and cerebro-spinal fluid (CSF) voxel masks. Finally, functional images were spatially smoothed with an isotropic 8-mm full-width-half-maximum Gaussian kernel and high-pass filtering with a 128-s cut-off to reduce low-frequency drift.

### Physiological artefact correction

During the 7T-fMRI acquisition of fMRI experiment 1, we recorded the cardiac pulse signal and respiration of participants by means of an MRI compatible pulse oximeter and pneumatic belt (Siemens Healthcare, Erlangen, Germany) at a sampling rate of 50 Hz. In addition, scanner pulses were recorded in an analogue input for synchronisation of fMRI and physiological data at 200 Hz. Due to technical problems, these data were not available for all scanning blocks and participants (average of 2.7 blocks, range 0 to 5 blocks per participant). Physiological artefact correction was performed for fMRI data with available concurrent physiological data. This involved band-pass filtering the pulse data between 20 and 150 bpm (0.3 and 2.5 Hz, respectively) to improve peak detection. Subsequently, RETROICOR was used to create regressors that were fed into the subject-specific fMRI analyses (GLMs) as confound regressors to remove spurious fluctuations. Fluctuations due to cardiac and respiratory phase were each modeled by 6 regressors for sine and cosine Fourier series components extending to the 3rd harmonic. Two additional regressors modeled lower frequency changes in respiration and heart rate with a sliding window analysis following ^51^.

### Region-of-interest (ROI) definition

Based on our a priori hypothesis ^13, 19^, ROI analyses were performed for the right entorhinal cortex (EC). Right EC ROIs were created on the Montreal Neurological Institute (MNI152) T1 template using a probabilistic atlas based on cytoarchitectonic mapping of ten human post-mortem brains ^52^ with FSL 5.0.4 (http://fsl.fmrib.ox.ac.uk/fsl/). The probability threshold was conservative (95%) for the estimation of hexadirectional orientations and liberal (0%, i.e. including all voxels with non-zero probability) for the small volume correction of the mask. Thresholded masks were binarised and converted to NiFTI file format and then normalised to the space of the individual functional images via the group-specific EPI template (Figure 5 – figure supplement 1) using the Advanced Neuroimaging Toolbox (ANTS; http://www.picsl.upenn.edu/ANTS/). SPM was used to reslice the ROI mask dimensions to the EPI dimension, which was again followed by binarisation of the masks. Through the same procedure, a right, primary visual cortex ^53^ mask (95% threshold) and a mamillary body ^54^ mask (25% threshold) were created for control analyses.

### Analysis of fMRI time series

Following pre-processing, fMRI time series were modeled with general linear models (GLMs). The different trial phases of the object-location memory task were modeled with two regressors. One regressor was used for the retrieval phase (replacement of an object) and one for the encoding phase (following the location response, when the object was shown at the correct location and could be collected), both of which were associated with a parametric modulator for spatial memory performance to discount large-error trials. Inter-trial-intervals (presentation of a fixation cross on a grey background) were not explicitly modeled and served as an implicit baseline. The presentation of the object cues and the feedback was modeled with two additional regressors. Furthermore, all GLMs included nuisance regressors, comprising at least 6 movement parameters, 2 regressors for signal fluctuations across white and grey matter voxels and 1 regressor to model time points with frame-wise displacements ^55^ larger than 0.5 mm. In addition, physiological signals have been recorded for a sub-set of participants (see section below for details) which was used to correct for cardiac and respiratory artefacts by means of 14 additional regressors. The main regressors of interest modeled virtual movement periods with two associated parametric modulators (see ‘Analysis of hexadirectional activity’ below for details). Coefficients for each regressor were estimated for each participant using maximum likelihood estimates to account for serial correlations. All parametric modulators were normalized to have zero mean and thus be orthogonal to the un-modulated regressor. Prior to the second-level random effects analysis, the linear contrast images of the regression coefficients underwent nonlinear normalization to the group-specific template brain using ANTS.

### Analysis of grid-cell-like representations

The orientation of 6-fold rotational symmetry of entorhinal activity (referred to as ‘hexadirectional activity’ and consistent with grid-cell representations in humans^19^) was estimated in participant’s right EC using a quadrature-filter approach on fMRI data during fast movements in all trial phases. Participant’s virtual-navigation fMRI data entered a general linear model (GLM) with two parametric modulators of a movement regressor. These modelled the sine and cosine of running direction θ(t) in periods of 60° (i.e. sin(6*θ(t)) and cos(6*θ(t))) for participant’s 50% fastest movement time points, where grid-cell-like representations can be reliably detected ^13, 19^. Multiplication by 6 transformed running direction into 60° periodic space to create regressors sensitive to activation showing a six-fold rotational symmetry in running direction. Activations with six evenly spaced peaks as a function of running direction will produce parameter estimates β_1_ and β_2_ for the two regressors with large amplitude sqrt (β_1_² + β_2_²). To this end, running direction θ(t) was arbitrarily aligned to 0° of the coordinate system underlying the virtual reality engine. Participants were not aware of the environmental coordinate system. The relationship between the underlying coordinate system and the polarisation axes (defined by extra-maze cues) differed between fMRI experiment 1 and fMRI experiment 2. The orientation of the polarisation axis (i.e. 0°) had an angular offset from the underlying coordinate system of 15° in fMRI experiment 1 and 90° in fMRI experiment 2. This made it unlikely that an anchoring of grid-cell representations to polarisation axes were due to other factors, such as viewing direction during the start of the experiment, which was −15° in fMRI experiment 1 and −90° in fMRI experiment 2, relative to the visible polarisation axes. Next, the parameter estimates of the two parametric modulators (β_1_ and β_2_) were extracted from the right EC ROI and used to calculate preferred orientation in 60° space (varying between 0° and 59°). A participant’s mean orientation of hexadirectional activity was defined as φ_60°_ = arctan(β_1_/β_2_), where β_1_ is the averaged beta value for sin[6*θ(t)] and β_2_ is the averaged beta value for cos[6*θ(t)] across voxels of the right EC. Dividing by six transformed the mean orientation φ_60°_ back into standard circular space of 360° for one of the three putative grid axes (the remaining two being 60° and 120° relative to the first).

Our main research question was if environmental geometry affects the orientation of putative grid-cell representations (see below for a description of the statistical test procedure). To additionally cross-validate effects of entorhinal hexadirectional activity ^13, 14, 19^, we tested the temporal stability of preferred orientations and their regional specificity in a split-half procedure (Experiment 1). This was done only for experiment 1, because data acquisition was roughly twice as long, participants completed more trials (179 versus 117, on average; two-sample t-test, T_(48)_=5.25, p<0.001) and SNR likely higher due to high-field scanning compared to experiment 2, which warranted a sacrifice in sensitivity for the main research question.

The procedure involved testing activation differences in the second half of the data with six-fold rotational symmetry that was aligned with the (potentially environmentally determined) hexadirectional activity estimated from the first half of the data. More specifically, the second GLM contained regressors for both ‘aligned’ and ‘misaligned’ runs relative to the estimated hexadirectional activity (respectively, this means running directions were either less than ± 15° or more than ± 15° oriented relative to the nearest axis of hexadirectional activity). As for the estimation procedure, regressors modeling six-fold rotational symmetry captured participant’s 50% fastest movement time points. Participants’ contrast values (aligned > misaligned) then entered a second level random-effects analysis to test for hexadirectional activity in the entire brain volume acquired. Supra-threshold activation in the right EC would indicate temporal stability and regional specificity of putative grid orientation.

Having evaluated temporal stability and regional specificity of the quadrature-filter approach for investigation of grid-cell-like representations in fMRI experiment 1, we decided to maximise statistical power addressing the main research question of environmental effects on hexadirectional activity in fMRI experiment 2.

### Analysis of environmental anchoring of hexadirectional activity

We tested environmental anchoring of the hexadirectional activity relative to the polarisation axes by using a V test for circular homogeneity ^56^. The V test for circular homogeneity is similar to the Rayleigh test for circular homogeneity and can be used if an a-priori hypothesis of a certain mean direction in a sample of angles is being tested. Due to our hypothesis of a relationship between the orientation of the grid-system and anisotropy in spatial information derived from angular changes to polarising cues, we tested participant’s putative grid orientations in 0°-60° space (Figure 5B and Figure 7A) for the presence of a mean direction aligned 30° off the polarisation axis.

## Acknowledgements

CFD’s, MN’s and TNS’s research is supported by the Max Planck Society; the European Research Council (ERC-CoG GEOCOG 724836); the Kavli Foundation, the Centre of Excellence scheme of the Research Council of Norway – Centre for Neural Computation, The Egil and Pauline Braathen and Fred Kavli Centre for Cortical Microcircuits, the National Infrastructure scheme of the Research Council of Norway – NORBRAIN; and the Netherlands Organisation for Scientific Research (NWO-Vidi 452-12-009; NWO-Gravitation 024-001-006; NWO-MaGW 406-14-114; NWO-MaGW 406-15-291).

CB’s, NB’s and BWT’s research is supported by the Wellcome Trust. NB’s research is also supported by the European Research Council. The authors would like to thank A Backus, J Bellmund, P Medendorp and B Milivojevic for useful discussions, A. Vicente-Grabovetsky for help with data analyses, B Somai for help with optimising the behavioural paradigm and C Hutton for support with the physiological noise correction.

## SUPPLEMENTAL INFORMATION

### Supplemental Figures

**Figure 1 – figure supplement 1.**
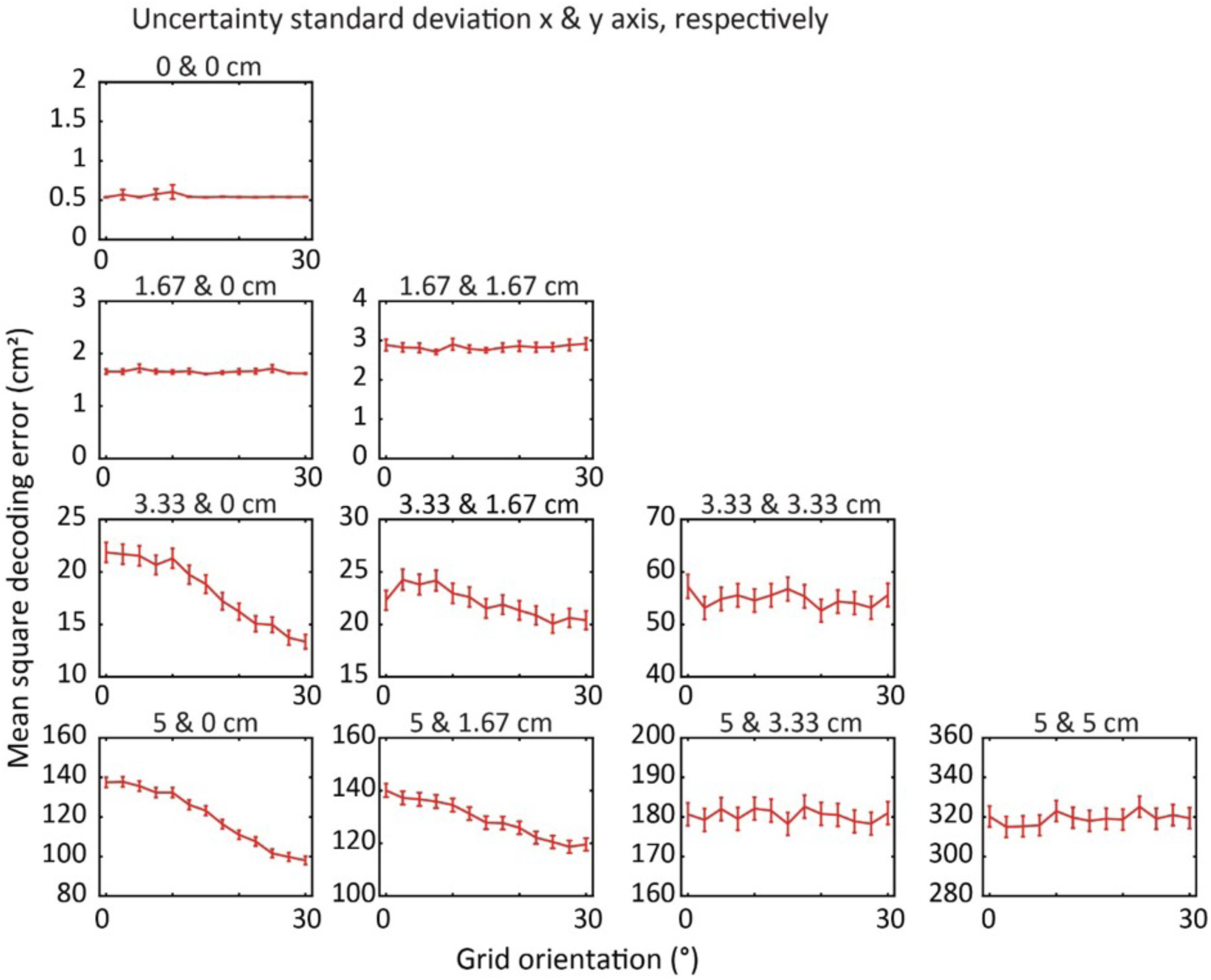
More extreme anisotropy in spatial uncertainty results in a more pronounced dependency of self-localisation accuracy on grid orientation. The performance of grid cell systems was assessed while independently varying the degrees of spatial uncertainty in two orthogonal axes. When uncertainty is equal in both axes performance does not depend on the orientation of the grid pattern. As uncertainty becomes more anisotropic, self-localisation is more accurate in grid cell systems in which the grid pattern axes are aligned away from the axis of greatest spatial uncertainty.

**Figure 1 – figure supplement 2.**
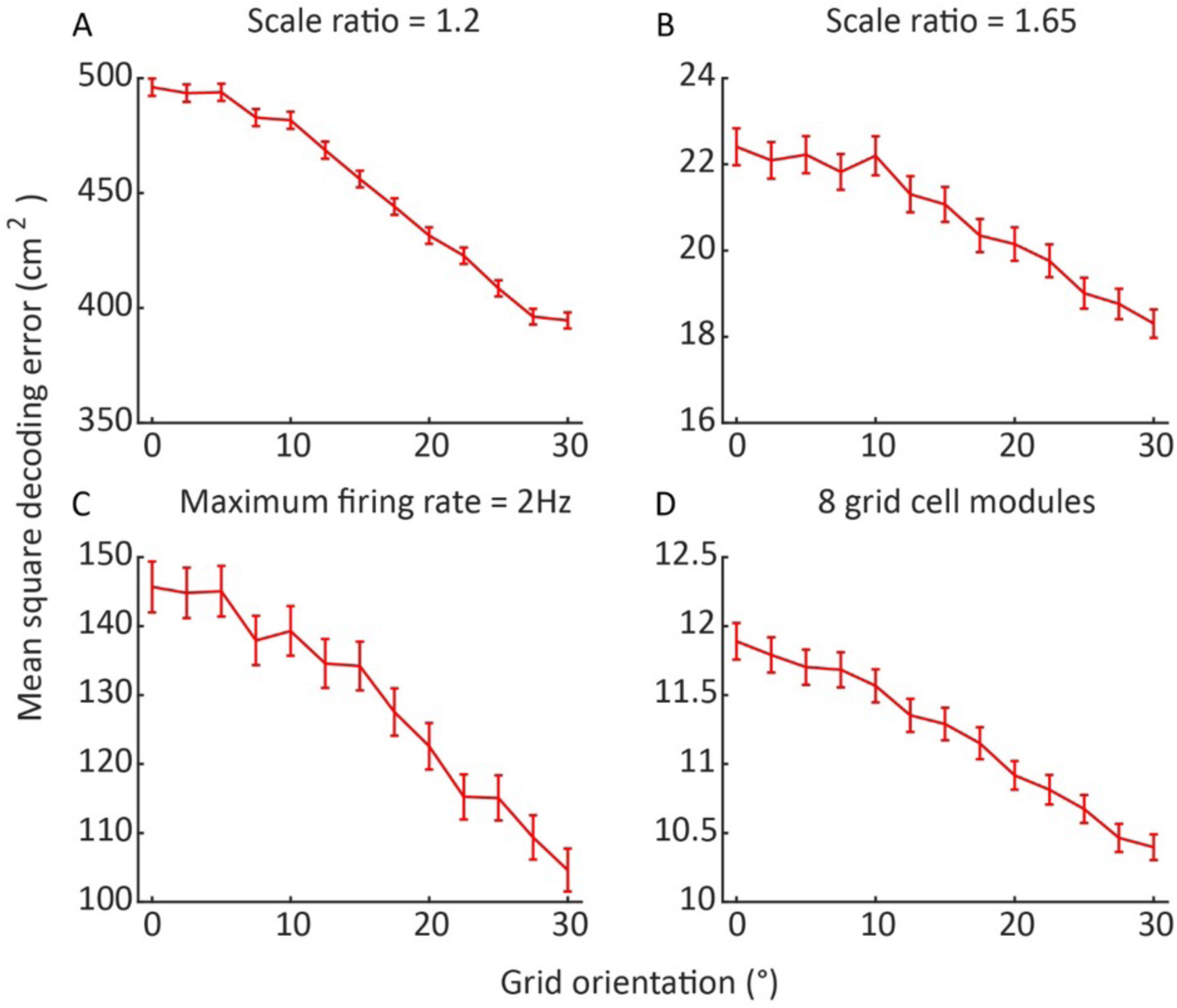
The dependency of self-localisation accuracy on grid orientation is stable within reasonable sets of grid cell system parameters. Performance is consistently best when the grid axes are aligned away from the axis of least spatial uncertainty, across variations in the parameters of the grid cell system. Error bars indicate 95% confidence interval, n = 150,000. **A** Grid period scaling factor reduced to 1.2. **B** Grid period scaling factor increased to 1.65**. C** Grid cell maximum firing rate reduced to 2Hz. (In order to compensate for increased effects of noise in this system, the number of cells per module was quadrupled. Due to the high computational intensity of this simulation n = 75,000.) **D** Four further grid cell modules added, with scales continuing to increase geometrically.

**Figure 1 – figure supplement 3.**
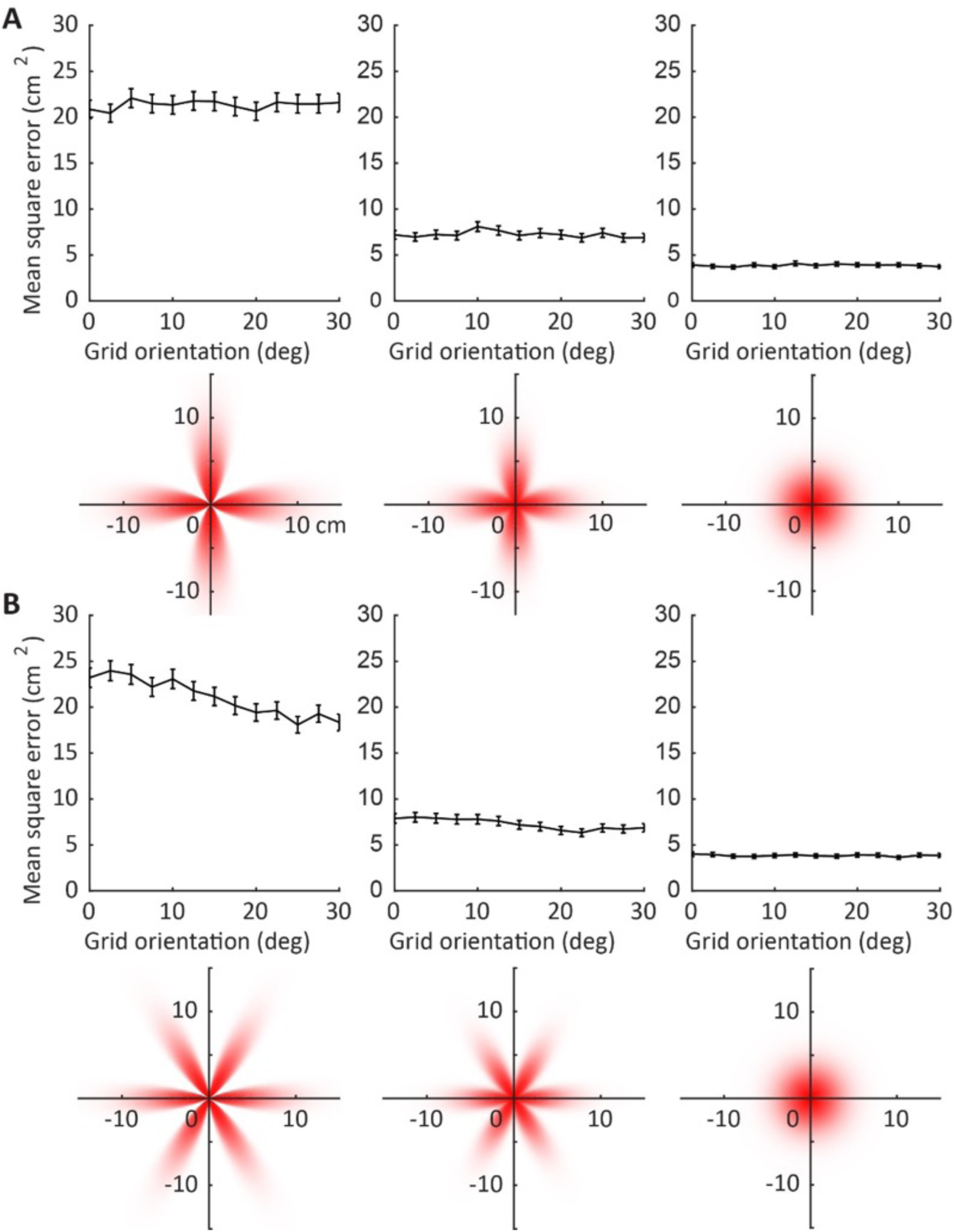
Higher-order uncertainty distributions and self-localisation accuracy. **A** Four-leaf uncertainty distribution. Across different grid orientations the different grid axes are variously aligned and misaligned with the directions of greatest and least certainty. No clear trend for an optimal grid orientation is apparent. **B** Six-leaf distribution. Across different grid orientations the grid axes are either aligned or misaligned with the high-uncertainty directions, producing the same trend as the simple (two-leaf) anisotropic uncertainty distribution. Localisation performance is best when the grid axes are aligned away from the axis of least spatial uncertainty. Error bars indicate 95% confidence interval, n = 150,000.

**Figure 1 – figure supplement 4.**
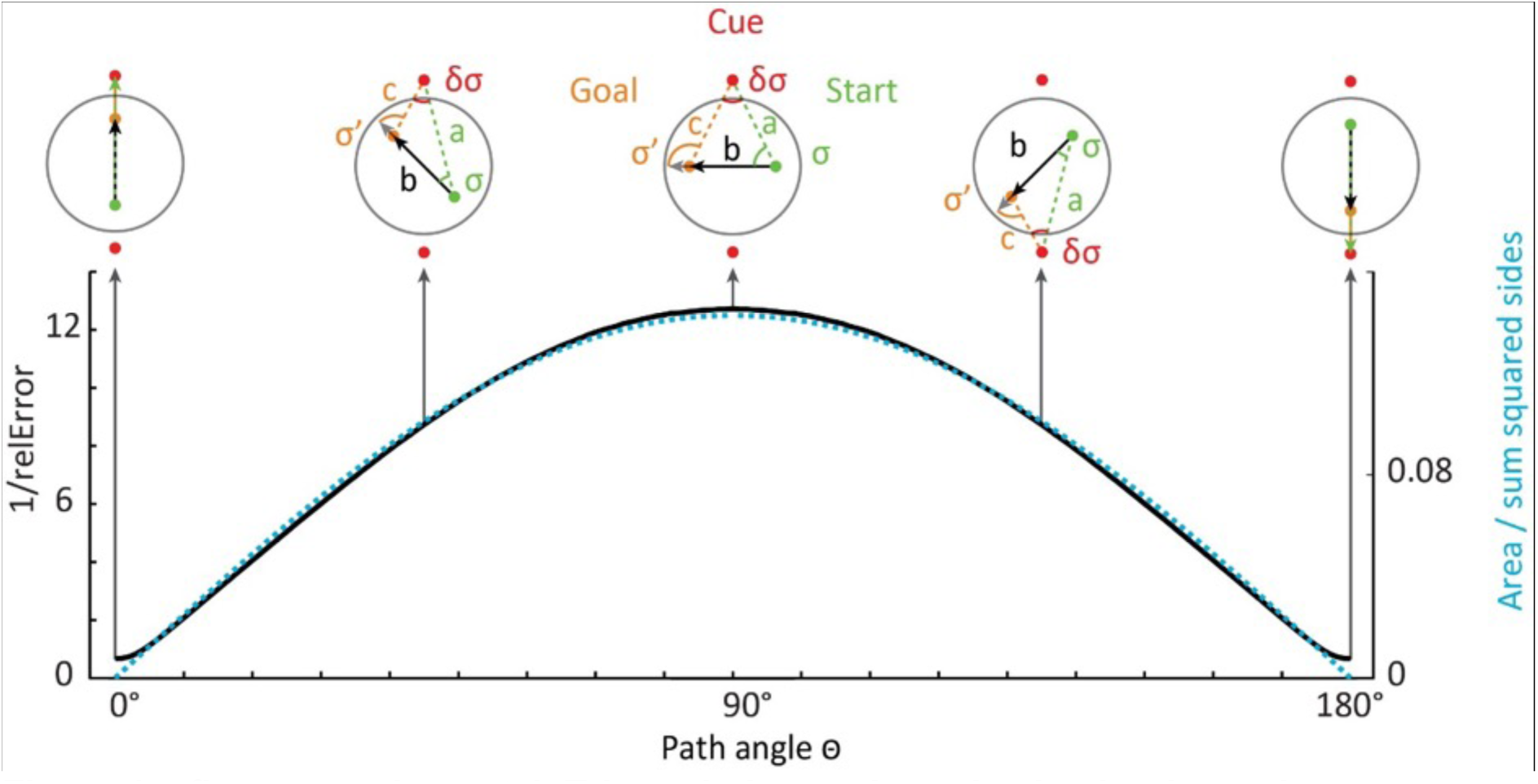
Triangulation under noise in circular and square environments. To test the impact of anisotropic optic flow information on spatial computations, we performed a biologically inspired simulation of Euclidean triangulation. For example, an estimate of the distance between start and end points was computed from noisy estimates of the angles and distance to one of the cues using equation 2. All sides (a-c) of a triangle served as both inputs and distance to be estimated, before the results were averaged on one iteration. The median noise resilience (1/relative error [relError]) across iterations is plotted in black. On a given iteration, relErr is determined as the absolute distance error / side length, averaged across the three sides of each triangle. Black arrows indicate example paths between two observer positions (start in green and goal in orange, always crossing the centre; see Materials and Methods for details). Red dots show polarising cues. Most accurate triangulation was achieved on paths orthogonal to the polarisation axis (10*10^3^ repetitions for each triangle, 90° ±15° versus 0° ±15°, two-sided Wilcoxon signed-rank test: Z=1026.42, p<0.001). Optimal path angle was well predicted by a quality measure for triangulation (triangle area / sum of squares of the side lengths; R=0.99, p<0.001). This measure increases for more equilateral triangles.

**Figure 1 – figure supplement 5.**
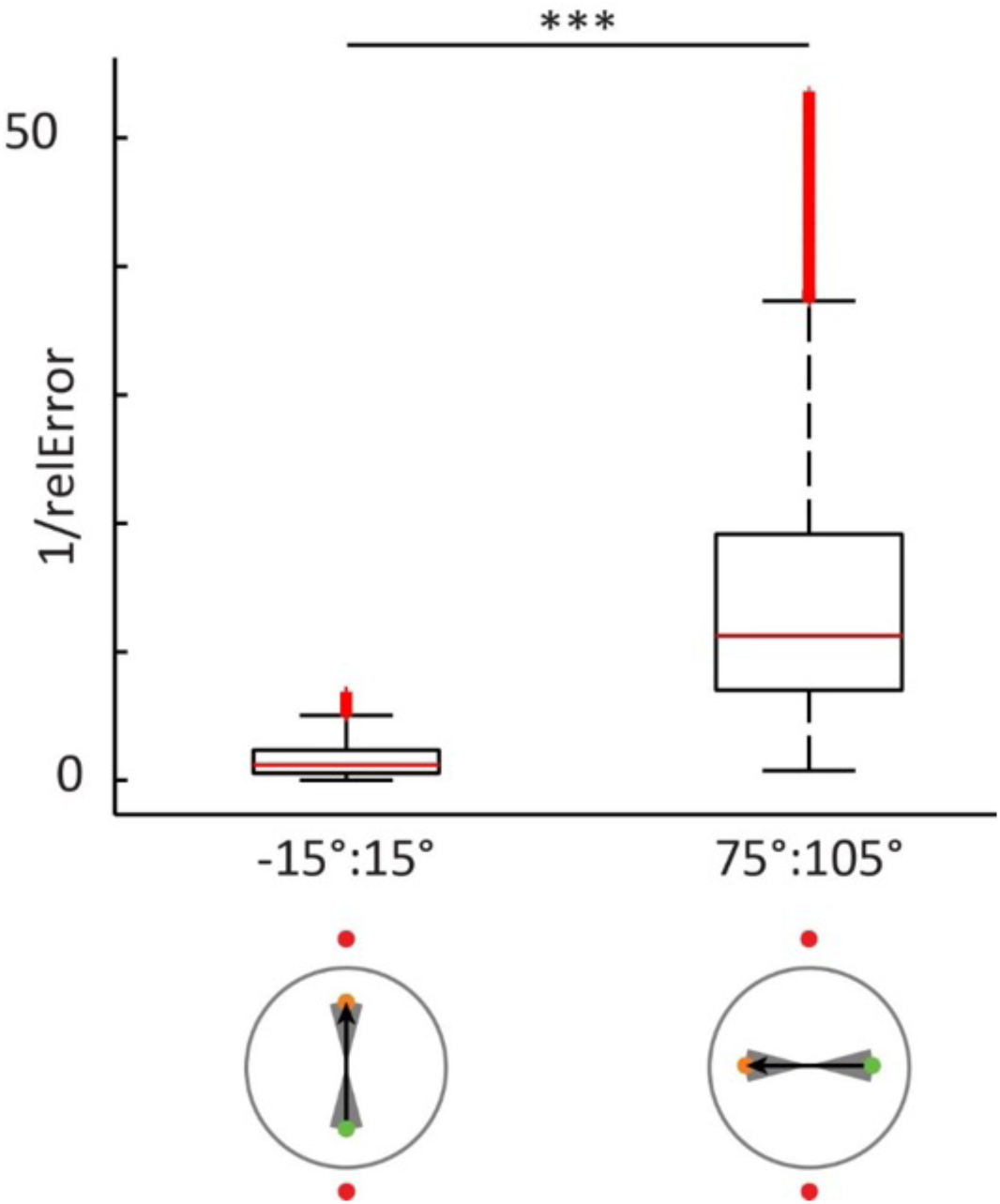
Magnitude of noise resilience of directional bin of paths centred on peaks and troughs of. **Figure S1**. The most accurate triangulation was achieved on paths orthogonal to the polarisation axis. 10*10^3^ repetitions for each triangle, 90° ±15° versus 0° ± 15°, two-sided Wilcoxon signed-rank test: Z=1026.42, p<0.001. The grey shaded area in the bottom panels indicate the range of paths that were tested. The box edges denote the 25th and 75th percentiles and central red mark the median. The whiskers extend maximally to *q*_3_ + *1.5 \** (*q*_3_ – *q*_1_) and minimally to *q*_1_ – 1.5 *\** (*q*_3_ – *q*_1_), where *q*_1_ and *q*_3_ are the 25th and 75th percentiles, respectively. A red + denotes points outside this range, with the exception of the upper 10% of values that were omitted for display purposes. Statistical testing included all data.

**Figure 1 – figure supplement 6.**
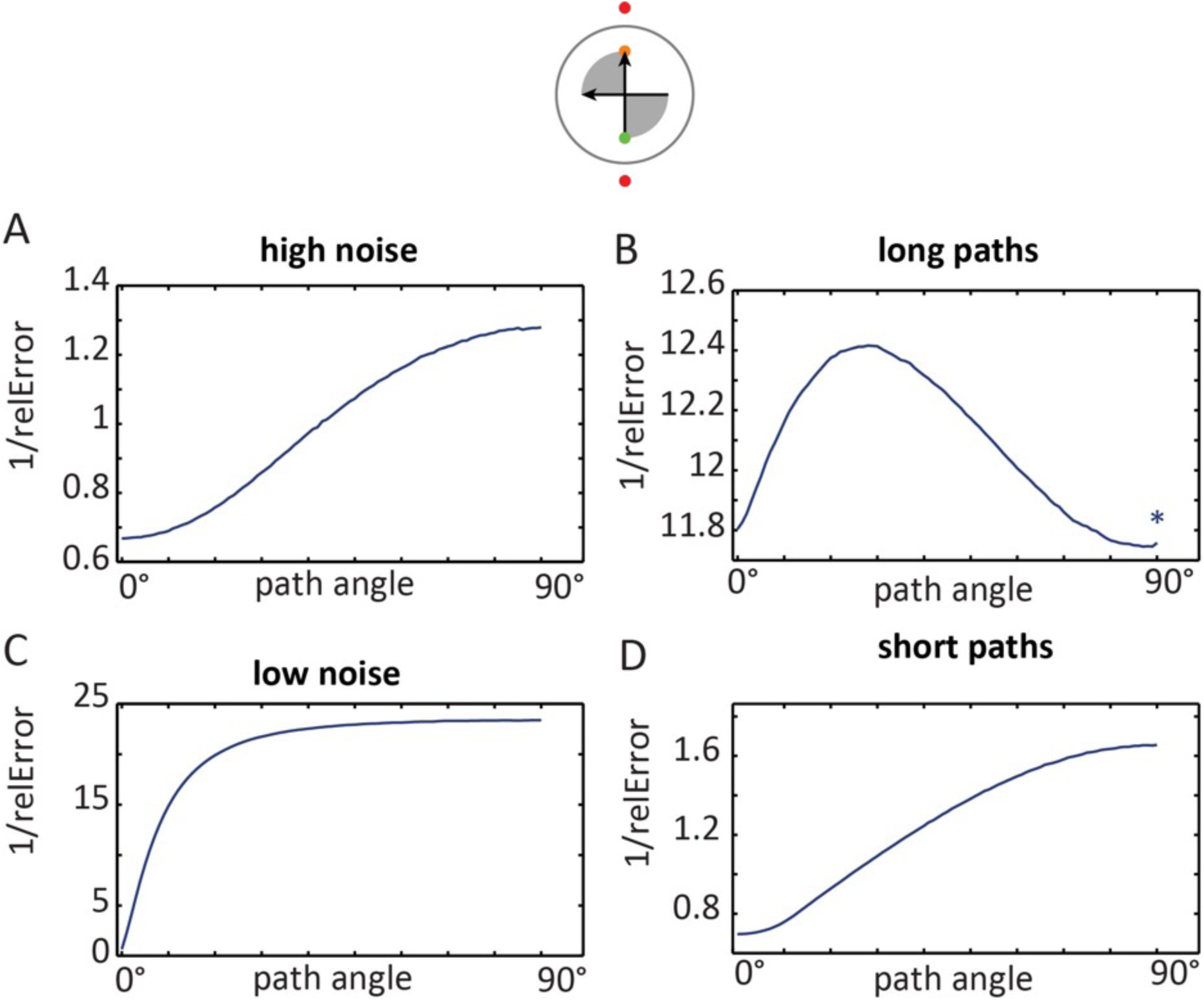
Effects of noise levels and path length on optimal triangulation paths. Environment with a single axis are defined by two cues (analogous to the two fMRI experiments and the behavioural experiment). All (A, C, D) except the long path condition (B) yielded an optimum at 90°. However, this extreme case never occurred for participant’s paths in the fMRI experiments due to the limitations of the circular boundary. The average length of straight (+-45°) paths was 11% of the polarisation axis’ length in fMRI experiment 1 (12% in fMRI experiment 2) – See Figure 3 – figure supplement 1E-F. High noise = randomly sampled from a distribution with a 10 times larger sigma (62.6, instead of 6.26, see Materials and Methods). Low noise = randomly sampled from a distribution with a 10 times smaller sigma (0.626, instead of 6.26). Long paths = simulated path length was equal to the length of the polarisation axis (instead of 50%, see Materials and Methods). Short paths = simulated path length was 5% of the length of the polarisation axis (instead of 50%, see Materials and Methods). Asterix = plots have been smoothed with a 5°-wide kernel for display purposes. The grey shaded area in the top panels indicates the range of paths that were used.

**Figure 2 – figure supplement 1.**
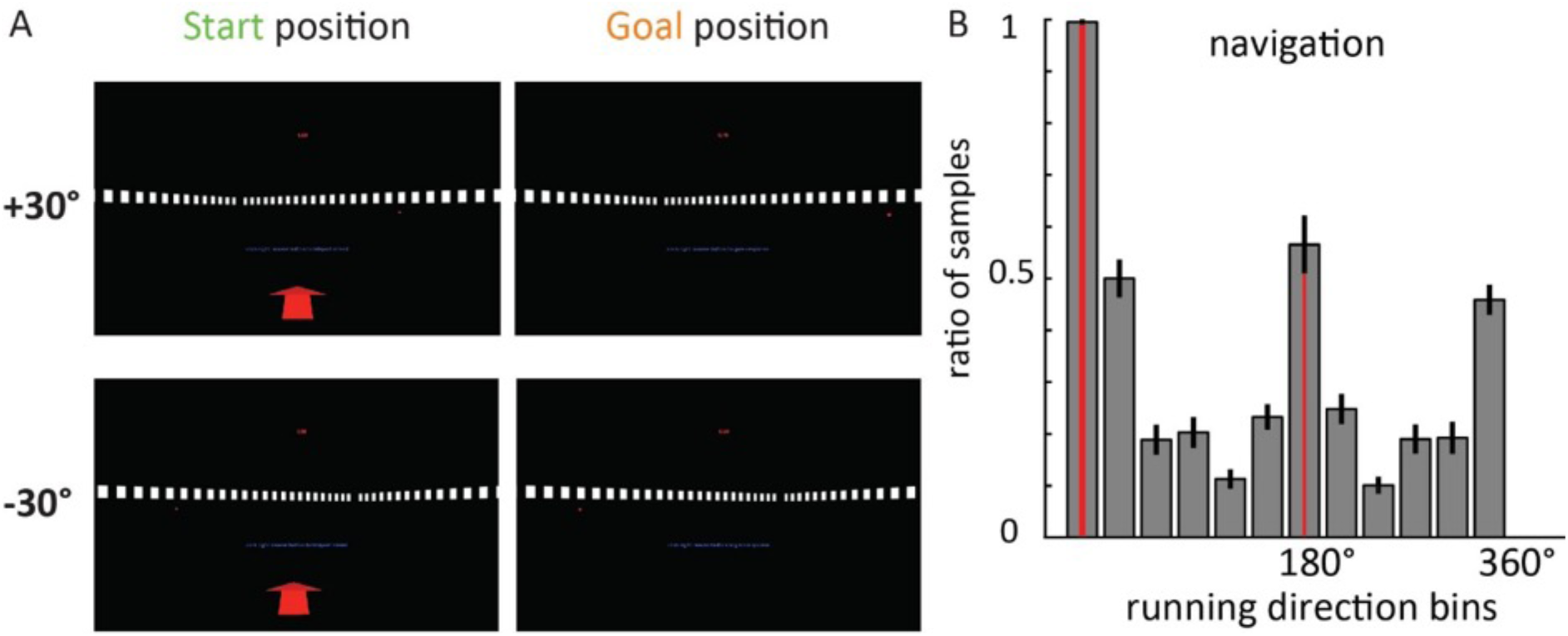
**A** Additional views of the behavioural experiment at the beginning (Start) and end (Goal) of a path. Note that the background was rendered at infinity (see Materials and Methods), such that it did not change during teleportation in the ±30° or the 0° condition (Figure 2BC). **B** Sampling of movement direction. Participants were not only exposed to translations in the ±30° and 0° condition, which might force spatial representations to align with those directions. During free navigation to a start position, all other directions were sampled, albeit not homogeneously (Friedman test: χ^2^_(11)_ =179, p<0.0001; note that on some trials the start position was on the opposite side of the environment).

**Figure 2 – figure supplement 2.**
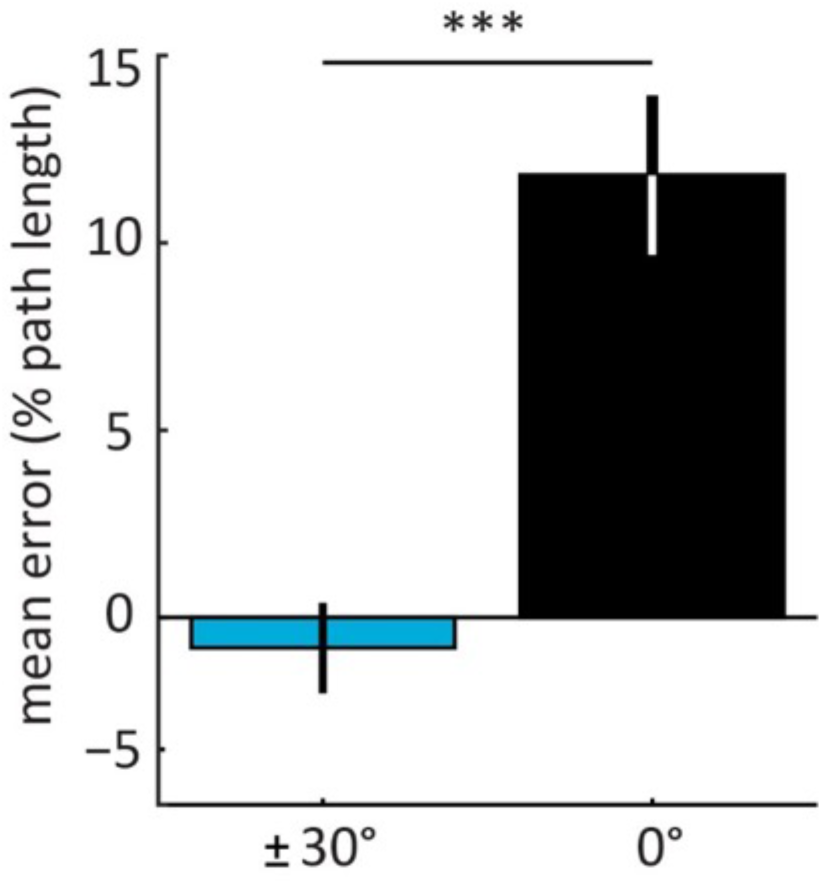
Mean distance estimation error (in percent of correct path length) in the behavioural experiment. (Figure 2). Paths along the polarisation axis (0°) yielded less accurate distance estimation than oblique paths (±30°). Paired, two-sided t-test N=20; T _(19)_ = 5.47, p<0.001. See Figure 2C for an effect in the same direction for absolute errors. Bars show errors in percent of correct path length averaged across participants +- S. E. M.; Mean 0° condition = 11.8 %; Mean ±30° condition= −0.8 %.

**Figure 3 – figure supplement 1.**
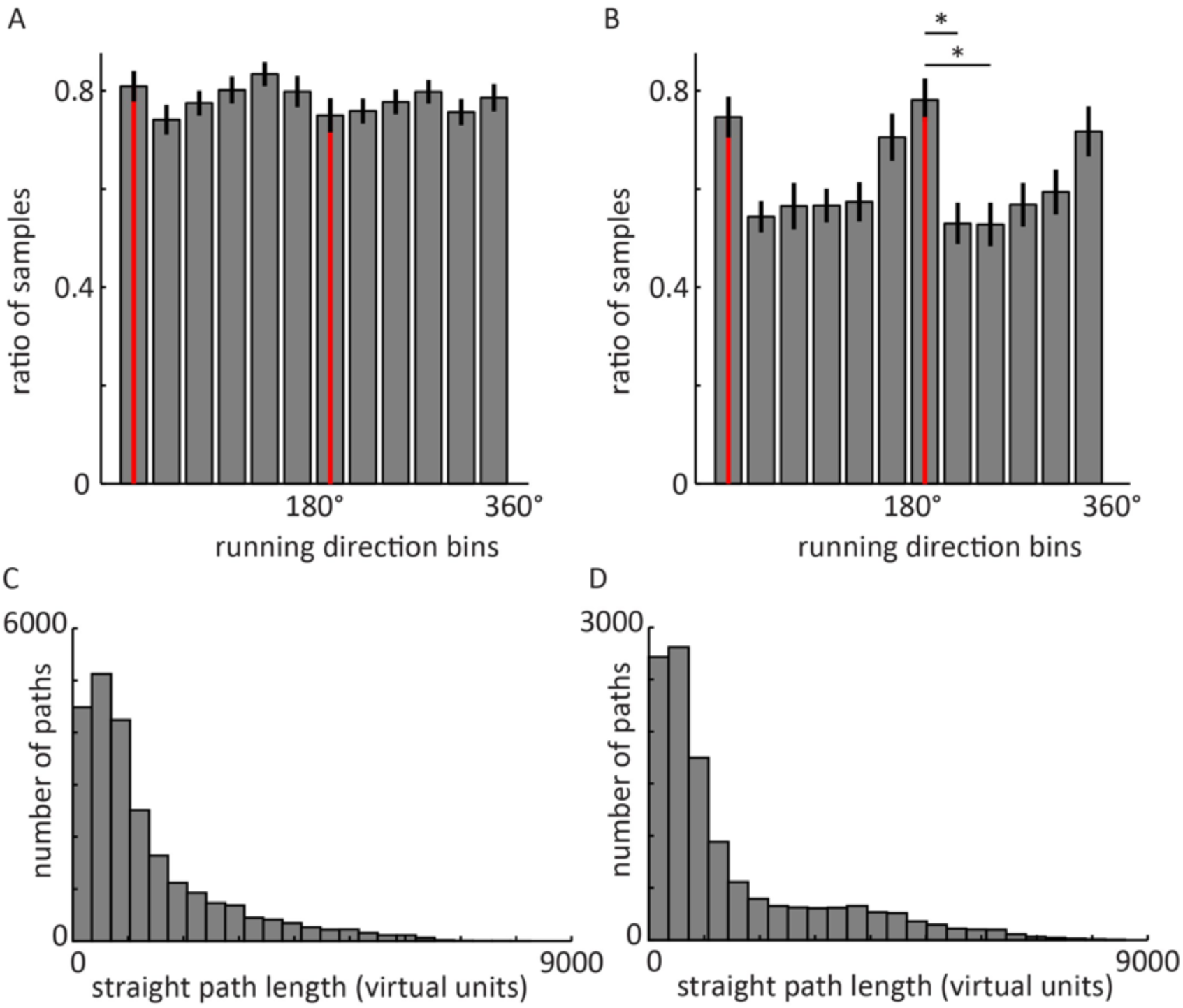
Behavioural analyses of fMRI experiment 1 (A, C, E) and fMRI Experiment 2 (B, D, F). Spatial memory performance: the decreases in drop error indicate that participants in both experiments were able to successfully navigate and remember locations in the sparse virtual environments. Participants learned the locations of 6 objects in fMRI experiment 1 (A) and 4 objects in fMRI experiment 2 (B; See Materials and Methods). Red line denotes mean drop error (i.e. Euclidean distance in virtual units between participants’ response location and the correct location of a given object on a given trial) across participants. Grey outline denotes standard error of the mean. For display purposes, results are shown up to trial number 90 for consistency. Variations in the number of trials across participants were due to differences in self-paced completion of trials. **C-D** Sampling of running directions. The number of samples of movements in 30° bins of running direction was normalised within participants for comparability across participants by dividing it by the maximum number of samples in any of the 12 bins, thereby yielding a maximum value of 1 for a bin. **C** FMRI experiment 1 (polarisation axis defined by configural cues), N = 26: A non-parametric Friedman test of median differences among repeated measures of directional sampling was revealed no clear differences (χ^2^_(11)_ =9.5, p=0.57). **D** FMRI experiment 2 (polarisation axis defined by non-configural cues), N = 24: A non-parametric Friedman test of median differences among repeated measures revealed differences (χ^2^_(11)_ =36.7, p<0.001). Post-Hoc tests with Tukey-Kramer correction for multiple comparisons revealed that particularly runs along the environmental axis at 165° (+-15°) occurred more often than runs oblique at 195° (+-15°) and 125° (+-15°). Asterix: p<0.05 Note that both the absence of a difference in fMRI experiment 1, as well as more frequent runs along the polarisation axis in fMRI experiment 2 speak against the possibility that the environmental effects on hexadirectional activity reported above would be due to biases in navigation behaviour. Error bars show S.E.M. over participants. **E-F** Distances of running paths. Histograms show the number of straight paths for different distances. Path length was determined as the Euclidean distance between start and end point of a path with continuous movement and rotations of less than +-45° (i.e. a 90°-wide bin). FMRI experiment 1 (E): mean = 1316.2 vu. FMRI experiment 2 (F): mean = 1493.5 vu.

**Figure 5 – figure supplement 1.**
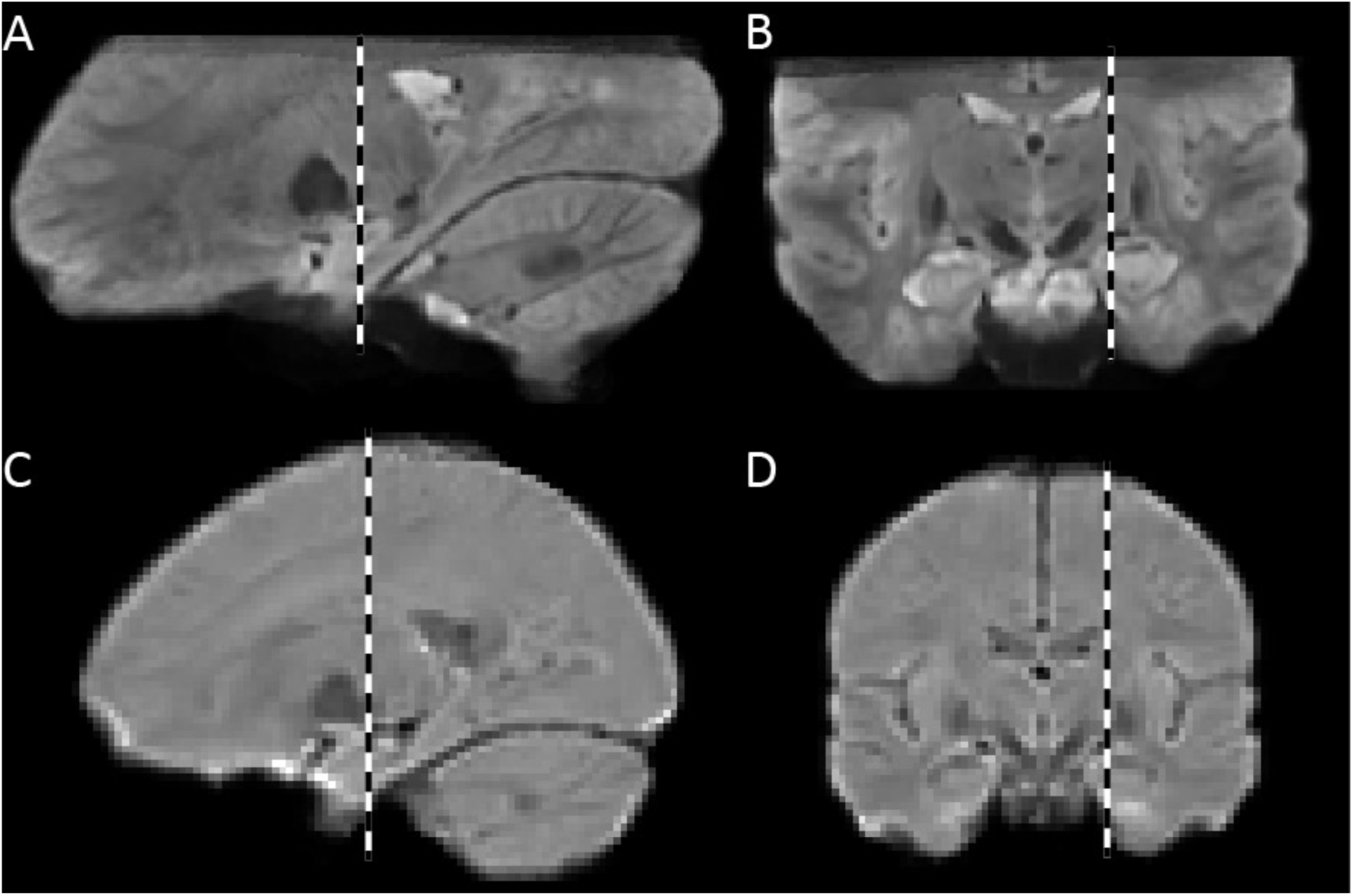
Mean functional images across participants used as template brains. A-B. Template for fMRI experiment 1 (7T scanner). **C-D** Template for fMRI experiment 2 (3T scanner). Dashed lines indicate the location of the slice in the corresponding orientation in the panel above or below. Template images were created with Advanced Neuroimaging Toolbox (ANTS; http://www.picsl.upenn.edu/ANTS/) based on individual, mean 3D echo-planar images. Note the relatively high contrast for functional images in the 7T data, with clear grey and white matter intensity differences even in the medial temporal lobes.

